# Resolving cell cycle speed in one snapshot with a live-cell fluorescent reporter

**DOI:** 10.1101/494252

**Authors:** Anna E. Eastman, Xinyue Chen, Xiao Hu, Amaleah A. Hartman, Aria M. Pearlman Morales, Cindy Yang, Jun Lu, Hao Yuan Kueh, Shangqin Guo

## Abstract

Cell proliferation changes concomitantly with fate transitions during reprogramming, differentiation, regeneration, and oncogenesis. Methods to resolve cell cycle length heterogeneity in real-time are currently lacking. Here, we describe a genetically encoded fluorescent reporter that captures live cell cycle speed using a single measurement. This reporter is based on the color-changing Fluorescent Timer (FT) protein, which emits blue fluorescence when newly synthesized before maturing into a red fluorescent protein. We generated a mouse strain expressing an H2B-FT fusion reporter from a universally active locus, and demonstrate that faster-cycling cells can be distinguished from slower-cycling ones based on the intracellular fluorescence ratio between the FT’s blue and red states. Using this reporter, we reveal the native cell cycle speed distributions of fresh hematopoietic cells, and demonstrate its utility in analyzing cell proliferation in solid tissues. This system is broadly applicable for dissecting functional heterogeneity associated with cell cycle dynamics in complex tissues.

## Introduction

Cell cycle speed varies widely and undergoes dynamic changes during development and tissue homeostasis, linking characteristic cycling behavior with unique aspects of regenerative and developmental biology (Chen et al., 2015; Soufi and Dalton, 2016). Across most of the animal kingdom, the cleavage divisions initiating embryonic development follow well-defined rapid and synchronous mitotic cycles (O’Farrell et al., 2004). Cell cycles lengthen and become heterogeneous at the onset of gastrulation (Deneke et al., 2016; Newport and Kirschner, 1982). In mammals, a characteristically fast cell cycle is seen in embryonic stem cells (ESCs) derived from the inner cell mass (White, 2005). Transition out of pluripotency, both *in vitro* and *in vivo*, is coupled with dramatic restructuring and lengthening of the cell cycle (Calder et al., 2013; White, 2005). Post-development, highly regulated cell cycles are seen across many tissue types including blood (Orford and Scadden, 2008; Pietras et al., 2011), brain (Yoshikawa, 2000), intestine (van der Flier and Clevers, 2009), and others (Liu et al., 2005; Tumbar et al., 2004). In tissues like the heart where cellular turnover is low, cells’ inability to re-enter the cell cycle appears to underlie the tissue’s poor regenerative capacity (Tzahor and Poss, 2017).

Abnormal cell cycle is characteristic of certain disease states such as cancer. Many oncogenes and tumor suppressor genes, such as Rb, p53 and c-Myc (Chen, 2016; Gabay et al., 2014; Knudsen and Wang, 2010), converge on the (dys)regulation of the cell cycle. Conventional chemotherapies often attempt to blunt cancer growth by targeting the cell cycle (Hamilton and Infante, 2016; Schwartz and Shah, 2005), but the efficacy could be compromised by natural heterogeneity in the proliferative behavior of the cancer cells (Fisher et al., 2013). Relapse due to development of chemo-resistance is thought to be related to the presence of quiescent cancer cells at the time of treatment (Chen et al., 2016). Recently, cyclin D-CDK4 has been shown to destabilize PD-L1 to induce tumor immune surveillance escape (Zhang et al., 2017).

Cell cycle control is equally important in non-malignant cell proliferation. In adult mice, it is estimated that most hematopoietic stem cells (HSCs) divide rarely (Wilson et al., 2008), and the ability to maintain quiescence is essential for their function (Pietras et al., 2011). Contrastingly, committed myeloid progenitors proliferate rapidly even under homeostasis (Passegue et al., 2005). Granulocyte-macrophage progenitors (GMPs) in particular appear to be one of the most proliferative cell types (Passegue et al., 2005), and are known to possess unique cell fate plasticity beyond the hematopoietic fate (Guo et al., 2014; Ye et al., 2015).

Overall, understanding the consequences of diverse cycling behaviors in development, regeneration, and disease is of fundamental importance. However, convenient assessment of cell cycle speed, especially in live cells of complex tissues, remains technically challenging.

Currently available strategies for cell cycle analysis have several limitations. First, they mostly focus on specific cell cycle phases (Bajar et al., 2016; Hagting et al., 1999; Sakaue-Sawano et al., 2008), but not on how quickly a cell progresses from one mitosis to the next. While a fast-dividing cell population usually contains a greater proportion of S/G2/M cells at any given time, any single cell could be transiting through these phases irrespective of its cell cycle speed. In fact, the presence of higher proportions of S/G2/M cells could indicate cell cycle arrest at these phases. Second, although image tracking of consecutive mitoses is direct and accurate for determining cell cycle length, many cells *in vivo* are not amenable to direct imaging due to their deep location and/or highly migratory behavior. Additionally, at least two mitoses need to occur for microscopy-based cell cycle quantification. Further, microscopy-based identification does not enable physical separation of fast-vs. slow-cycling cells for downstream molecular or functional assays. Third, while various label retention techniques have yielded much of our current knowledge on stem cell quiescence *in vivo* (Falkowska-Hansen et al., 2010; Tumbar et al., 2004; Wilson et al., 2008), the cycling kinetics of dividing cells are difficult to resolve after they lose their label during the chase period. Label retention assays, either in the form of generic dyes (Lyons et al., 2001), nucleotide analogs (Wilson et al., 2008), or genetically encoded H2B-GFP (Falkowska-Hansen et al., 2010; Tumbar et al., 2004; Wilson et al., 2008), reflect the cell divisional history during the chase period, but give little information about the current cycling state. The resolution of such methods is also limited by how similar the cells are to each other (Guo et al., 2014). Some of the labels are also cytotoxic, or require fixation to visualize (Wilson et al., 2008). To overcome the above-mentioned limitations, we implemented a genetically encoded color-changing fluorescent protein which reports on cell cycle speed in a ratiometric manner. This reporter is suitable for cell cycle studies of live cells both *in vitro* and *in vivo*.

## Results

### Mathematical modeling predicts that long- and short-lived molecules are differentially susceptible to molecular halving by cell division

The steady state level of a given molecule is determined by the sum of its biogenesis and turnover. Molecular turnover occurs by active catalysis or by passive dilution when cells divide. The extent to which cell division contributes to molecular turnover depends on a given molecule’s half-life (Equation S1). When the time scale of turnover is significantly shorter than the length of the cell cycle, the contribution from active turnover dominates. An example of this is the inhibitor of NF-κB, or IκBa, which undergoes enhanced degradation upon phosphorylation within minutes of signaling, leading to NF-κB activation (Hacker and Karin, 2006). However, when turnover is slow and approaches the length of the cell cycle, molecular dilution by cell division becomes the main mode of removal. This phenomenon is seen during myeloid fate commitment from multipotent hematopoietic progenitors, where the stable PU.1 protein accumulates as the cell cycle lengthens (Kueh et al., 2013). Therefore, the intracellular concentration of a stable molecule varies more than that of an unstable molecule in response to cell division rate. The different behaviors of short-vs. long-lived molecules in relation to cell cycle length are illustrated in Figure 1A, suggesting a potential strategy for ranking cell cycle length by measuring two types of molecules, e.g. fluorescent proteins, that are distinct in their half-lives.

**Figure 1.**
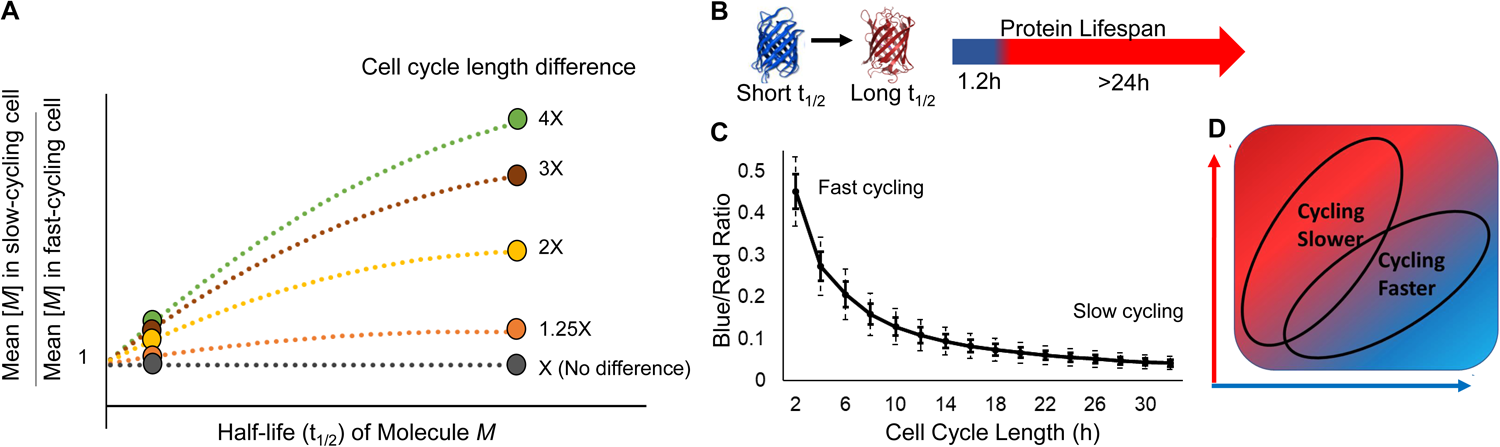
Design principles of a fluorescent reporter of cell cycle speed. **A**. Based on Equation S1, intracellular levels of a given molecule, *M*, are predicted to depend on its molecular half-life as well as the cell cycle lengths. Here, the ratio of M’s concentration in theoretical cell populations with different cycling speeds are plotted. When *M* is short-lived (circle, far left), its cellular concentration shows little difference between slow- and fast-cycling cells. However, with a long-lived *M* (circle, far right), the concentration difference increases proportionally to the difference in cell cycle lengths. **B.** The color-changing Fluorescent Timer (“FT”) displays a short half-life as a blue protein, and a long half-life as a red protein. **C.** The average blue/red ratio of cells expressing the FT is predicted to drop as cell cycle lengthens (see Equation S2), providing a fluorescence-based strategy to identify cells cycling at different rates. Because the blue/red ratio fluctuates when cell cycle progresses (see Figure S1B-D), the relationship between blue/red ratio and cell cycle length is best described by a probability distribution. For modeling, all cells are assumed to maintain a constant cycling rate through generations. Solid and dashed error bars denote 1 and 2 standard deviations, respectively. All FT blue/red ratio are defined as blue / (blue + red). **D.** The anticipated positions of slow and fast cycling cells on a hypothetical plot of blue and red fluorescence intensity.

To exploit the differential susceptibility of labile and stable molecular species to cell division-mediated halving, we turned our attention to the Fluorescent Timer (FT) (Subach et al., 2009). The FT is a color-changing protein that emits blue fluorescence when newly synthesized, and irreversibly turns red after a characteristic time delay (Subach et al., 2009) (Figure 1B, Figure S1A). The blue and red proteins are synthesized orthogonally in a stoichiometric 1:1 manner, because the blue species is a folding intermediate of the red, whose abundance can be measured separately by fluorescence microscopy and/or by flow cytometry. Importantly, the immature blue form of the FT is lost quickly after synthesis when it changes into the red form, which has a much longer half-life (Figure 1B, Figure S1A). Thus, the blue form models the short-lived molecule, whose concentration is less variable across different cell cycle rates as compared to the stable red form (Figure S1B-D). From a mathematical model describing the kinetics of this fluorescence conversion and subsequent degradation, we find that the steady-state blue/red fluorescence ratio scales inversely with cell cycle length (Equation S2), with a shorter (longer) cell cycle length producing a higher (lower) blue/red ratio. Thus, the blue/red FT ratio decreases as the cell cycle lengthens or slows (Figure 1C, Figure S1B-D), providing a potential strategy to distinguish cycling rates (Figure 1D).

### H2B-FT fusion proteins undergo blue to red conversion, and are suitable for proliferative cells

To render the FT useful for microscopy-based quantitation, the FT was fused to the C-terminus of histone H2B (Figure 2A), a widely used chromatin-tagging strategy with minimal adverse consequences (Hadjantonakis and Papaioannou, 2004). This localized the FT signal to the chromatin (Figure 2B), facilitating nuclear segmentation and image processing. To assess whether tagging the FT onto a histone interferes with its color-changing properties, we imaged Hela cells expressing the H2B-FT under the control of a Tetracycline-inducible (TetO) promoter (Figure 2A). HeLa cells transduced with the “Medium” FT variant (Subach et al., 2009) (Figure S1A) began to produce detectible blue signal ∼2 hours after adding doxycycline (Dox) to the culture medium (Figure 2C-D, Movie S1), a time delay presumably required for Dox-induced transcription and translation. Approximately 1.5 hours after the first appearance of blue fluorescence, red signal emerged as the first wave of FT molecules underwent maturation (Figure 2C, Movie S1). The average level of red fluorescence per nucleus continued to increase over the next 26 hours until reaching steady-state fluorescence (Figure 2E), i.e. the level of fluorescence in cells continuously maintained in Dox. By contrast, the average level of blue fluorescence plateaued much sooner (after ∼7 hours) (Figure 2D), as expected for a molecule which is lost (in this case converted to red) shortly after synthesis. To confirm the divergent turnover times for the blue and red FT, we treated cells with Dox briefly (“Dox Pulse”, Figure 2F). Upon Dox washout, blue signal dropped below detectible levels within 7 hours. In contrast, the red signal persisted for the remainder of the imaging experiment, >24 hours (Figure 2E-F), confirming that the red H2B-FT is more stable than the blue form. Mathematical simulation of the FT behavior predicts that it reports cell cycle speed only after its expression reaches steady-state (Figure S1B-D). When expression is first induced, the blue form should be over-represented independently of cell cycle effects because insufficient time has elapsed for the red molecules to accumulate. In agreement with the model, cells displayed higher blue/red ratio within the first 24 hours of Dox induction as compared to cells maintained continuously in Dox (Figure 2G, Movies S1 and S2). As such, all subsequent experiments described below were conducted after the cells had been exposed to Dox for sufficient time (≥2 days) to reach steady-state H2B-FT expression.

**Figure 2.**
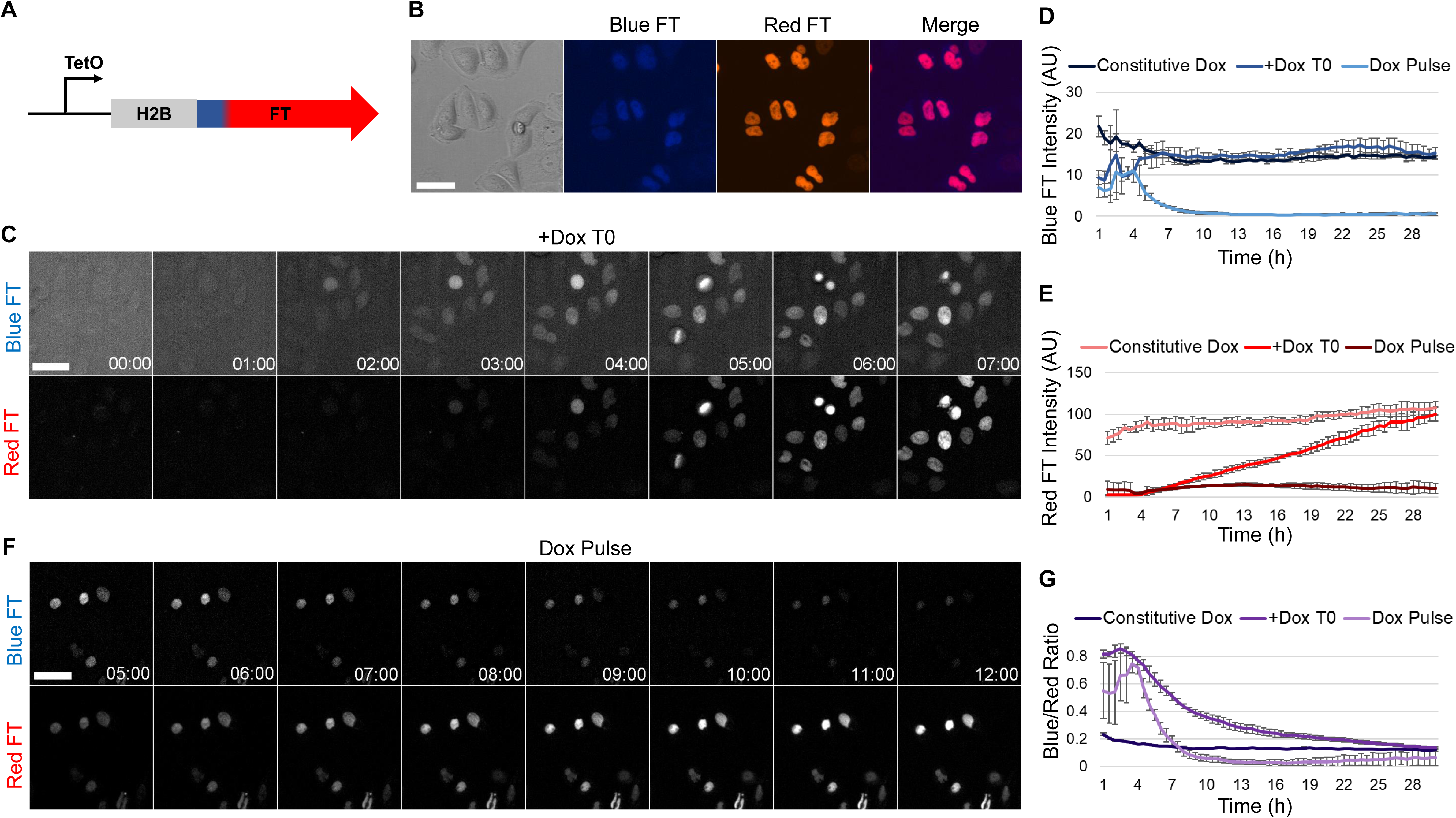
Characterization of the H2B-FT reporter in cultured cells. **A**. Schematic of inducible H2B-FT expression construct. **B.** Representative images of HeLa cells expressing the H2B-FT. **C.** Selected time series of H2B-FT fluorescence in HeLa cells from a representative field of view following doxycycline (Dox) induction. **D-E**. Quantification of average nuclear blue (D) and red (E) FT intensity over time following Dox treatment. **F.** Selected time series of HeLa cells after Dox washout at T=4.5 h. **G**. Data from (D-E) plotted as blue/red ratio. Error bars represent standard deviation across n = 3 culture wells. Nuclear intensity measurements were normalized to timepoint 0. Scale bars = 50µm.

We next assessed the reduction in H2B-FT-red signal contributed by cell division-dependent and cell division-independent processes. We tracked single cells (primary mouse embryonic fibroblasts (MEFs) or Hela) over time after Dox washout (Figure S2A-B). In MEFs, the red fluorescence decreased marginally in non-dividing MEFs (irradiated or naturally non-dividing by chance) (Figure S2C). In contrast, mitosis led to a ∼50% drop in red fluorescence (Figure S2A,C). Additionally, we tracked the combined red fluorescence intensity from all progeny arising from single Hela cells by mitosis (Figure S2B,D). Similar to MEFs, mitosis caused ∼50% drop in red fluorescence, as long as all the progeny were accounted for (Figure S2D). These analyses confirmed that the predominant turnover mode of H2B-FT-red is mediated by cell division. Of note, the half-life of H2B-FT-red (∼84 hours in MEFs and ∼37 hours in Hela) appears to be longer than the 24-hour half-life reported for the unfused FT-red (Subach et al., 2009). The increased H2B-FT stability could result from being fused to a core histone, which are among the most stable proteins (Toyama et al., 2013). These results indicate that the resolution of the H2B-FT as a cell cycle speed reporter could become limited in slow-dividing cells, where non-division mediated decay would be expected to influence the levels of red protein and consequently the blue/red ratio.

### H2B-FT blue/red ratio responds to cell cycle manipulation in cultured cells

The mathematical model predicts that cell cycle acceleration causes the blue/red ratio to increase, while cell cycle lengthening leads to the opposite effect (Figure 1C-D, Figure S1C-D). To test this experimentally, we perturbed the cell cycle by multiple methods in cultured cells expressing the H2B-FT.

We first accelerated cell proliferation by c-Myc expression in primary MEFs carrying allelic expression of the H2B-FT (details below), specifically the FT-Medium variant (Figure S1A). Following overnight transduction with c-Myc or an empty vector (EV) control virus, the H2B-FT fluorescence was determined by quantitative time-lapse microscopy (Figure S3A-B). As expected, cells overexpressing c-Myc proliferated more rapidly (Figure 3A). On the population level, c-Myc-transduced MEFs displayed a higher blue/red ratio than control MEFs after 70 hours but not at 25 hours post-transduction (Figure 3B), which was presumably before the exogenously expressed c-Myc had significantly altered the cell cycle. Notably, the blue/red ratio was heterogeneous at the single-cell level in both control and c-Myc-expressing cultures (Figure 3C). To determine whether this heterogeneity reflected cell cycle speed heterogeneity, we categorized cells based on their relative blue/red ratio following a FACS-style gating strategy (Figure 3C), and systematically determined individual cell cycle lengths by tracking the time interval between consecutive mitoses within the same cell lineage. In c-Myc transduced cultures, there were more cells in the high blue/red ratio group (“Blue cells”) and fewer cells of low blue/red ratio (“Red cells”) (Figure 3C-D). Importantly, the blue cells divided faster than the red cells in both conditions (Figure 3E). While most red cells had a long cell cycle (>30 hours), the blue cell cycle length centered around 10-15 hours/cycle in the control culture and 5-10 hours/cycle in the c-Myc-transduced culture (Figure 3E, blue cells). A small population (∼1.5%) of c-Myc-transduced cells had a cell cycle length shorter than 5 hours/cycle (Figure 3E). Interestingly, for most of the cells that had an intermediate level of blue/red ratio (cells between the gates in Figure 3C), their cell cycle length centered around 25 hours/cycle, likely representing the gap between the fast dividing blue cells and the slow dividing red cells (Figure 3E). These results are in good agreement with previous reports of MEF cell cycle length (White, 2005). Importantly, the blue/red ratio of individual cells reflected cell cycle rates regardless of whether they had been transduced with c-Myc. In either case, cells with the highest blue/red ratio divided faster (Figure 3E-F). As such, the blue/red ratio of H2B-FT reports cell proliferation rate at both the population and single-cell level.

**Figure 3.**
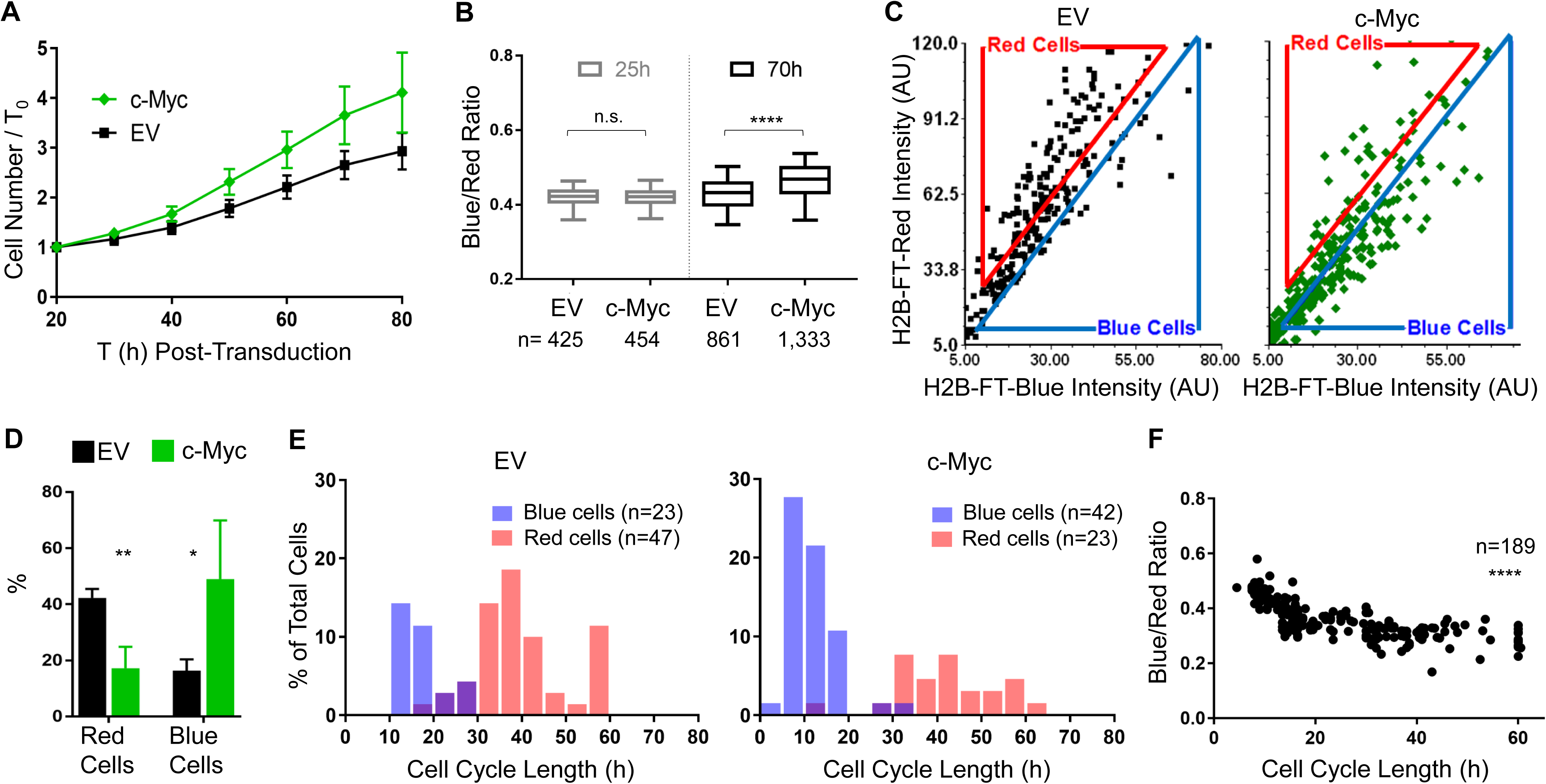
The H2B-FT blue/red profile reflects cell proliferation rate *in vitro*. **A.** Changes in cell number over time in primary iH2B-FT MEFs following transduction with either c-Myc or empty vector control (EV). Error bars denote standard deviation across n=4 culture wells. **B.** The ratio of blue/red fluorescence intensity was determined for individual cells at 25h and 70h post transduction. Box plots represent the median and interquartile range and whiskers represent 5th-95th percentile. P=0.295 (25h) and P<0.0001 (70h), determined using Mann-Whitney test with a 99% confidence interval. **C.** The blue and red fluorescence level of individual cells at 70h post transduction. Each dot denotes a single cell. FACS-style gates were applied to representative scatter plots. **D.** The percentage of cells within each gate. Error bars show standard deviation across n=4 culture wells. P=0.0035 (Red Cells) and P=0.0498 (Blue Cells), determined using Student’s T-Test with Welch’s correction, 95% confidence interval. dF=4.057 (Red Cells) and dF=3.221 (Blue Cells). **E.** The cell cycle lengths of individual cells from each gate were determined by image tracking. Cell cycle length represents the time interval between two consecutive mitoses within the same cell lineage. Cell cycle length is heterogeneous and the distribution of cell cycle lengths for each condition are shown as histograms. n values refer to the number of cells tracked for each condition. **F.** The relationship between cell cycle length and H2B-FT blue/red ratio. Each dot denotes an individual cell. All trackable cells (n=189) from both conditions are plotted. Spearman correlation coefficient = −0.7808. P<0.0001 was calculated with a 95% confidence interval.

To test whether cell cycle lengthening would decrease the blue/red ratio, we expressed the H2B-FT in BaF3 cells, whose proliferation is dependent on IL-3 (Ajjappala et al., 2009). To slow their proliferation, BaF3 cells initially cultured in full IL-3 (270pg/ml) were switched to low IL-3 levels (14pg/ml). Within 48 hours after switching, BaF3 cells showed reduced proliferation, which completely ceased in IL-3 free conditions (Figure 4A). The blue/red fluorescence intensity of BaF3 cells cultured in these conditions were measured by flow cytometry. This analysis revealed that BaF3 cells grown in high IL-3 largely emitted blue fluorescence, which shifted toward the red axis at low IL-3 and became entirely red with IL-3 withdrawal (Figure 4B, Figure S3B-C).

**Figure 4.**
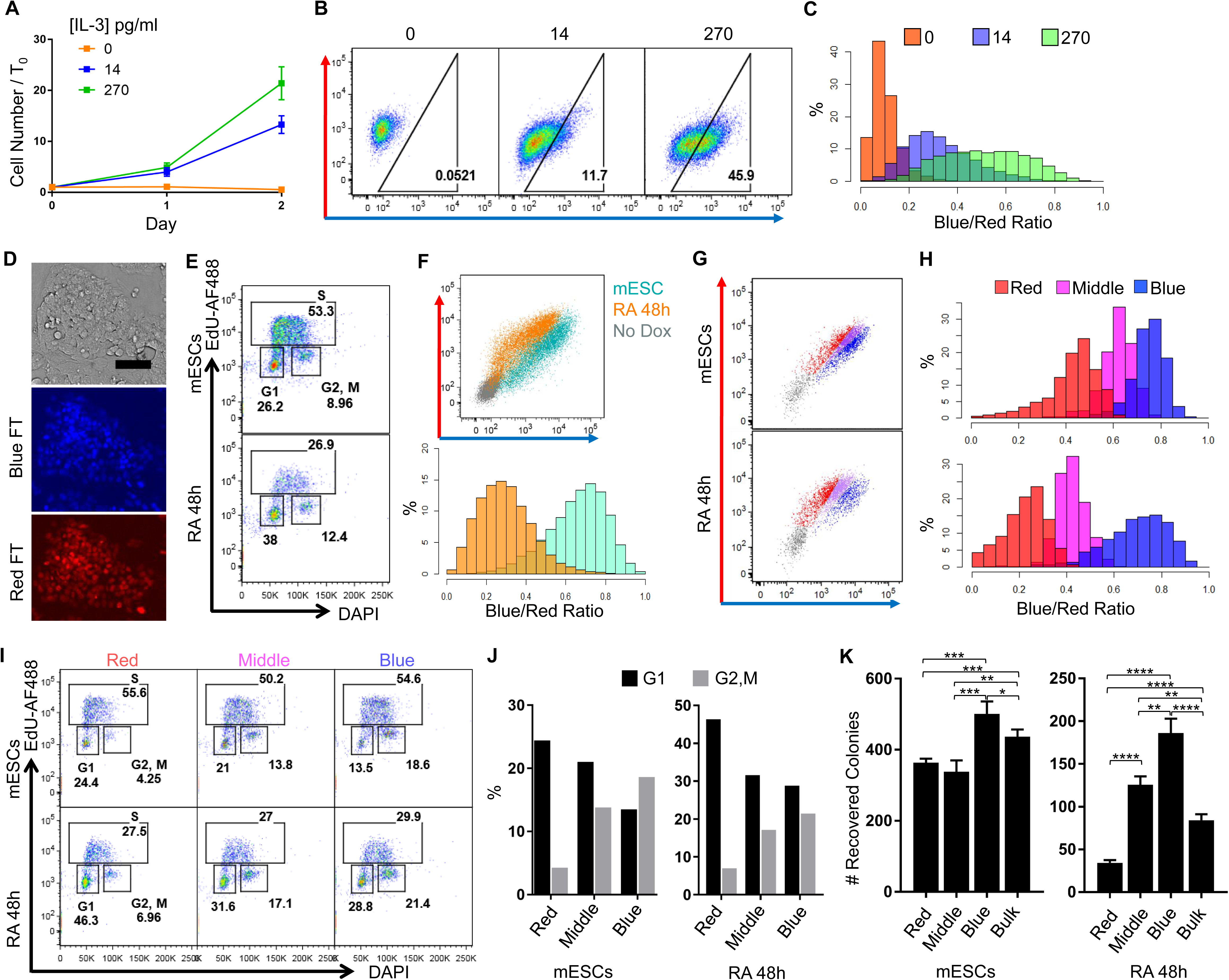
H2B-FT blue/red profile allows FACS sorting of live cells with different proliferative rates. **A.** BaF3 cells proliferate at different rates in varying IL-3 concentrations, as determined by cell counting. Error bars denote standard deviation across n=6 culture wells. **B.** Representative FACS plots of BaF3 cells expressing the H2B-FT reporter grown under different IL-3 concentrations. **C.** Histograms of blue/red fluorescence ratio derived from FACS data in (B). **D.** Representative colony morphology and FT fluorescence of H2B-FT-Medium knock-in mESCs maintained in feeder-free conditions. **E.** Confirmation of cell cycle change following 48h of RA treatment, as analyzed by EdU pulse-labeling and DNA content profiling. **F.** Representative FACS plots of red vs. blue fluorescence in pluripotent and RA-treated mESCs (top). These FACS data were re-plotted as histograms of blue/red ratio (bottom). **G-H.** Pluripotent and RA-treated mESCs stably transfected with H2B-FT via Sleeping Beauty transposon were FACS sorted according to their blue/red FT ratio. Representative gating strategy is shown. H2B-FT-negative gate was determined using non-transfected WT mESCs. **I.** DAPI/EdU cell cycle profiles of sub-populations sorted from mESCs and RA-treated cells. **J.** Frequency of cells in G1 vs. G2/M from the FACS-sorted populations shown in (I). **K.** The number of alkaline phosphatase positive colonies formed by the same number of cells sorted in (G) following 6 days of culture. Error bars denote standard deviation across n=3-4 culture wells. Significance determined using Student’s T-Test with a 95% confidence interval, df=6 (mESCs) and df=5-6 (RA 48h). Exact P values provided in Table S3. Scale bar in (D) = 80µm.

To facilitate direct comparison across large numbers of cells in FACS data, we used a normalized blue/red ratio, given by dividing a cell’s blue signal intensity by its combined blue and red intensity:

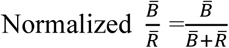

This normalization eliminates the difference in the amount of H2B-FT protein expressed by individual cells, and places all on a scale of 0-1, with 0 being all red and 1 being all blue (Figure 4C, Figure S4A-E). All graphs involving blue/red ratio were normalized in this manner. As expected, the blue/red ratio of BaF3 cells decreased in response to cell cycle lengthening (Figure 4C, Figure S3C-D). IL-3-induced differences in proliferation were captured by all three kinetic variants of the FT protein: H2B-FT-Fast, H2B-FT-Medium, and H2B-FT-Slow (Figure 4B-C, Figure S3C-D). Furthermore, as predicted by Equation S2, the range of blue/red ratio was lowest with FT-Fast and highest with FT-Slow (Figure S3C-D), which have the shortest and longest blue-to-red conversion time, respectively (Figure S1A). For simplicity, all data presented from here onward were obtained with the FT-Medium.

To examine the response of the H2B-FT reporter in a situation where cell cycle speed changes concomitantly with a cell fate transition, we assessed mouse embryonic stem cells (mESCs) with either allelic expression (Iacovino et al., 2011) (details below and Figure 4D), or transposon-mediated expression (Hudecek et al., 2017) of H2B-FT. mESCs have a characteristic cell cycle of 8-10 hours that immediately lengthens upon differentiation (White, 2005). Taking advantage of this well-defined change in cell cycle rate, we induced mESC differentiation with retinoic acid (RA) (Wichterle et al., 2002) (Figure S4F-G), which slowed the cell cycle as confirmed by EdU/DAPI staining (Figure 4E). As expected, mESCs treated with RA for 48 hours displayed a profound, population-wide shift toward the red-axis (Figure 4F). Only incomplete differentiation was expected, since the RA treatment was brief. This partial differentiation strategy yielded cell state heterogeneity, which was captured by the prominently red-shifted, but overlapping, population following RA treatment (Figure 4F). Taken together, our results demonstrate that the H2B-FT blue/red ratio responds to diverse modes of cell cycle manipulations in multiple cell types.

### H2B-FT blue/red profile enables FACS sorting of live cells with different cycling rates

We wished to test the feasibility of using the blue/red profile to physically separate live cell populations with distinct proliferation dynamics. To do this, mESCs kept in pluripotency maintenance conditions or treated with RA as described above were assessed in more detail. We sorted the cells based on their blue/red intensities (Figure 4G-H) and compared their cell cycle profile by DNA content staining (Figure 4I). As cells with longer cell cycles tend to increase their dwell time in G1 phase relative to other phases, we expected slow-dividing cells to be enriched for G1 cells. Consistently, the reddest population (Figure 4G-J, Red) was enriched for G1 phase cells, while the bluest (Figure 4G-J, Blue) contained a larger proportion of G2/M cells. Cells from the middle FT gate (Figure 4G-J, Middle) had an intermediate cell cycle profile. These results are consistent with the observations that pluripotent stem cells do not respond homogeneously or synchronously to differentiation induction (Drukker et al., 2012; Semrau et al., 2017). Cell cycle rate heterogeneity was not only seen in the RA-treated samples, but also for cells kept in pluripotency maintenance conditions (Figure 4G-J), corroborating reports that cellular heterogeneity exists under pluripotency maintenance conditions (Furusawa et al., 2006; Stewart et al., 2006) and is often associated with the cell cycle (Furusawa et al., 2006; Gonzales et al., 2015). These findings further validate that the H2B-FT blue/red ratio reports proliferation rate.

To evaluate functional heterogeneity associated with the H2B-FT color profile, cells from individual blue/red gates were FACS-sorted and re-plated in pluripotency conditions to evaluate their ability to give rise to new alkaline phosphatase (AP)-expressing colonies (Kalkan et al., 2017; Sela et al., 2012). We expected the slow-cycling (red) cells to be more differentiated and have reduced ability to form colonies, and that any cells which had remained pluripotent during the RA treatment to be among the fastest-cycling (blue) group. In concordance with the DAPI/EdU results, RA treatment greatly reduced the overall colony-forming activity (Figure 4K). Importantly, the residual colony-forming potential was most enriched in the blue cells and depleted in the red ones (Figure 4K, right). Our results also demonstrated that even among cells never induced to differentiate, colony forming potential was highest within the bluest cells, albeit this difference was less pronounced (Figure 4K, left). These results confirmed the presence of cell cycle heterogeneity in both culture conditions which could be captured and sorted by FACS using the H2B-FT reporter.

### The proliferative landscape of live hematopoietic cells as captured by the H2B-FT

To test whether the H2B-FT can report cell cycle speed in live cells *in vivo*, we took advantage of the well-known proliferative differences (Passegue et al., 2005) in hematopoietic stem and progenitor cells (HSPCs). For a first approach, the H2B-FT was virally introduced into donor HSCs, which were subsequently transplanted into recipient mice. Following engraftment and reconstitution, we analyzed the hematopoietic stem cells and multipotent progenitors (Lin-Kit+Sca+, LKS), the granulocyte-macrophage progenitors (GMPs), and the whole bone marrow (WBM) (Figure S5A-B, gating strategy). The heterogeneity as reflected by the blue/red profile was most exaggerated in the WBM (Figure 5A-B), consistent with the fact that WBM contains the entire dynamic spectrum from rapidly-cycling progenitors to post-mitotic differentiated cells. By contrast, the GMP and LKS compartments displayed narrower distributions (Figure 5A-B). Among the HSPCs positive for H2B-FT, LKS cells were red-shifted as compared to the GMPs (Figure 5A-B), in agreement with the LKS compartment containing the quiescent and slow-cycling hematopoietic progenitors, while the GMPs are highly proliferative (Passegue et al., 2005; Wilson et al., 2008). These reconstituted LKS cells were heterogeneous, containing a subset showing increased blue signal (Figure 5A-B), possibly reflecting the more proliferative multipotent progenitors (Passegue et al., 2005; Wilson et al., 2008). Although the GMPs were collectively bluer than LKS and many other cells, they were not the bluest within the bone marrow (Figure 5B). Further investigation revealed the bluest cells to be Ter119+ (Figure 5C-D), indicating erythroid identity (Kina et al., 2000). Contrastingly, Mac1+ myeloid cells displayed a wide distribution of blue/red ratio, with a severely red-shifted subpopulation (Figure 5C-D). Therefore, the proliferative behavior of hematopoietic cells, as established by BrdU uptake kinetics in earlier studies (Passegue et al., 2005; Pop et al., 2010), can be conveniently recapitulated by the H2B-FT expressed from a lentiviral vector.

**Figure 5.**
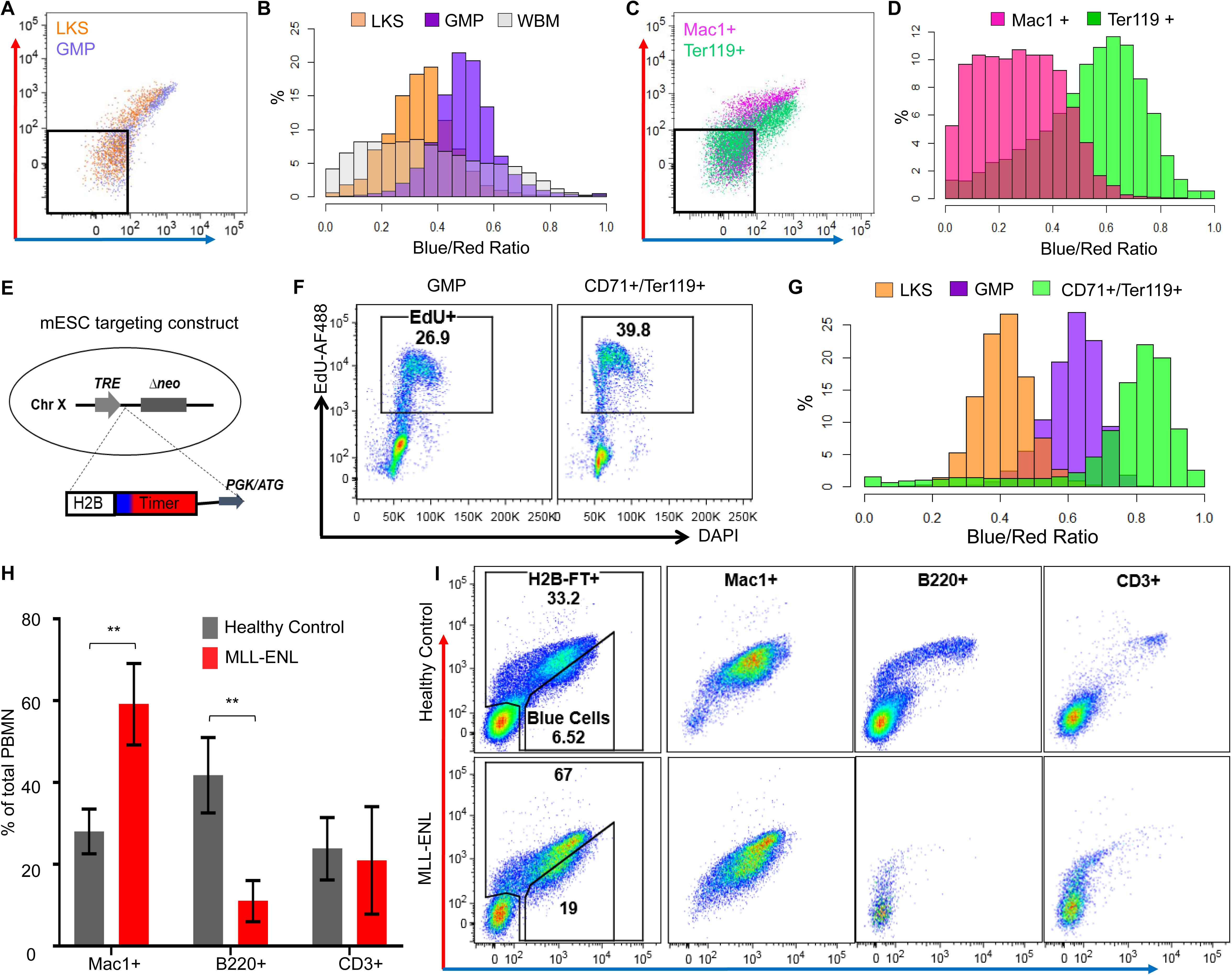
The proliferative landscape of live hematopoietic cells as captured by the H2B-FT reporter. **A.** Representative FACS plots of blue/red profile in LKS and GMP cells from reconstituted mice 2 months after transplantation with HSPCs virally expressing the H2B-FT reporter. **B.** Distribution of blue/red ratio in H2B-FT-expressing LKS and GMP populations, overlaid on that of whole bone marrow. **C.** FACS plots of myeloid (Mac1+) and erythroid (Ter119+) cells from reconstituted mouse bone marrow. **D.** Data from (C) plotted as histograms of blue/red H2B-FT ratio. Gates in (A) and (C) were used to exclude the non-transduced (H2B-FT-negative) cells. **E.** Targeting strategy for the *HPRT::*TetO-H2B-FT mouse allele. **F.** DAPI/EdU cell cycle profiles of GMPs vs. early erythroid cells following 2 hours of *in vivo* EdU labeling. **G.** Distributions of H2B-FT blue/red ratio in LKS, GMPs, and early erythroid cells. **H.** Frequency of myeloid (Mac1+), B-cell (B220+), and T-cell (CD3+) lineages in peripheral blood of healthy H2B-FT knock-in mice vs. those crossed with MLL-ENL (n=3 mice per group). All mice were treated with Dox for at least 8 days prior to analysis. Error bars represent standard deviation. P=0.0090 (Mac1+), P=0.0072 (B220+), and P=0.7629 (CD3+) determined using Student’s T-Test with a 95% confidence interval, dF=4. **I.** Representative H2B-FT blue/red profiles within defined peripheral blood subsets as in (H).

To eliminate the limitations associated with viral H2B-FT expression, we proceeded to generate a knock-in mouse by targeting the H2B-FT coding sequence into the *HPRT* locus (Iacovino et al., 2011), under the control of a Dox inducible promoter (*iH2B-FT*) (Figure 5E) to enable H2B-FT induction in desired cell types when crossed with specific rtTA or tTA. To test this mouse model and confirm the findings with the virally expressed H2B-FT, we crossed the *iH2B-FT* allele with Rosa26:rtTA (Hochedlinger et al., 2005), which should enable inducible H2B-FT expression in most tissues including the hematopoietic system. The compound mice were healthy and fertile. To directly compare the proliferative behavior of erythroid progenitors and GMPs, we pulsed the iH2B-FT mice with EdU for 2 hours and analyzed the cell cycle profile of CD71+/Ter119+ cells and GMPs (Figure S5A-C, gating strategy). Approximately 40% of the erythroid cells, as compared to 27% of GMPs, incorporated EdU during the labeling period (Figure 5F). Accordingly, the CD71+/Ter119+ cells were further blue-shifted as compared to GMPs (Figure 5G), corroborating the results obtained with the viral approach. When the HSPCs were analyzed, the LKS compartment was again robustly red-shifted and heterogeneous (Figure 5G), in agreement with the virally expressed H2B-FT (Figure 5A-B). Taken together, the known cycling rates of primary bone marrow cells are faithfully and reproducibly captured by the H2B-FT blue/red ratio irrespective of the expression strategy. The knock-in allele allows detection of physiological cell cycle rates without any further manipulation or labeling.

Circulating blood cells primarily consist of B cells (B220+), T cells (CD3+) and neutrophils/monocytes (Mac1+) (Figure 5H), with the myeloid cells turning over more rapidly than the lymphocytes in normal hematopoiesis (Passegue et al., 2005). As expected, such a difference is readily reflected by the H2B-FT blue/red profile: Mac1+ cells were bluer than B220+ or CD3+ cells, whose blue/red profiles were highly heterogeneous (Figure 5I, top). While many circulating B and T cells did not express detectible H2B-FT levels, the H2B-FT+ frequency among myeloid cells was >95% (Figure 5I, top), on par with H2B-FT+ frequency in the whole bone marrow. Importantly, the B and T cells which did show H2B-FT expression contained a severely red-shifted subpopulation which could represent long-lived memory cells (Tough and Sprent, 1995). Although the exact identity and function of these various cellular subsets remain to be determined, FACS based separation should provide a convenient approach for their further study.

Lastly, we explored the iH2B-FT blue/red profile in a disease state characterized by overt proliferation, such as acute myeloid leukemia (AML). To induce AML, we crossed the iH2B-FT mice with a Dox inducible MLL-ENL (iMLL-ENL) allele targeted into the *Col1a* locus (Ugale et al., 2014). Mice harboring both the iH2B-FT and iMLL-ENL were treated with Dox. As expected, the myeloid compartment expanded within 2 weeks, when the bone marrow contained a preponderance of L-GMPs (Figure S5D) and the blood became dominated by Mac1+ cells (Figure 5H-I, Figure S5E, bottom). The expanded myeloid compartment was immediately visible by the increased number of H2B-FT+ cells in the unstained whole blood (Figure 5I, bottom). Further, the bluest-shifted subpopulation became ∼3x larger than that in healthy control mice. Thus, the distinct proliferative behaviors of malignant hematopoietic cells can be readily detected by the H2B-FT blue/red ratio. Overall, the proliferative landscape of the hematopoietic tissue that has been well surveyed by other approaches, can be revealed in one simple FACS assay.

### H2B-FT blue/red ratio unveils the geography of cell proliferation *in situ*

Since cell proliferation is compartmentally organized in many solid tissues, we evaluated the ability of the H2B-FT reporter to resolve such borders in sections cut from various adult and embryonic organs. In mice pulsed with EdU for 35 minutes, we first examined a representative slow-turnover organ as well as a proliferative one. For the former category, we chose the kidney, which has an extremely low proliferation rate under homeostasis (Kusaba et al., 2014). For the latter, we selected the pylorus. Like other regions of the stomach, the pylorus is a glandular tube lined with epithelium supported by non-proliferative nerve, blood vessel, and muscle cells (Treuting et al., 2012). Gastric epithelial glands are organized into base, isthmus, and pit regions (Kim and Shivdasani, 2016), with the most rapidly-dividing progenitors located in the isthmus. As expected, the blue/red ratio of kidney cells was much lower overall than that of the gastric epithelial cells (Figure 6A-B, Figure S6, top and middle). However, the pylorus tissue was heterogeneous in its blue/red ratio, with a subset of cells appearing as red-shifted as the kidney (Figure 6B, top and middle). Image-quantified blue/red ratio was binned into five populations (P1-5, Figure 6B), rendered into a pseudo-colored heatmap, and applied back onto the original images (Figure 6C-D). Notably, cells at the base of gastric glands had a very low blue/red ratio, juxtaposed abruptly with a region of cells of increased blue/red ratio (Figure 6A,C, middle). The co-localization of this color boundary with a dense band of EdU+ nuclei confirmed that the blue/red ratio identified the border between the mostly quiescent base compartment and the transit-amplifying region. Glands that were fully captured longitudinally from base to pit furthermore showed a declining blue/red ratio past the most proliferative zone (Figure 6C-D, “Pylorus I”), as expected. Gastric glands captured in cross-section, representing assorted positions along the base-pit axis, additionally showed the pattern of co-localization between regions of high blue/red ratio and high density of EdU+ cells (Figure 6C-D, “Pylorus II”).

**Figure 6.**
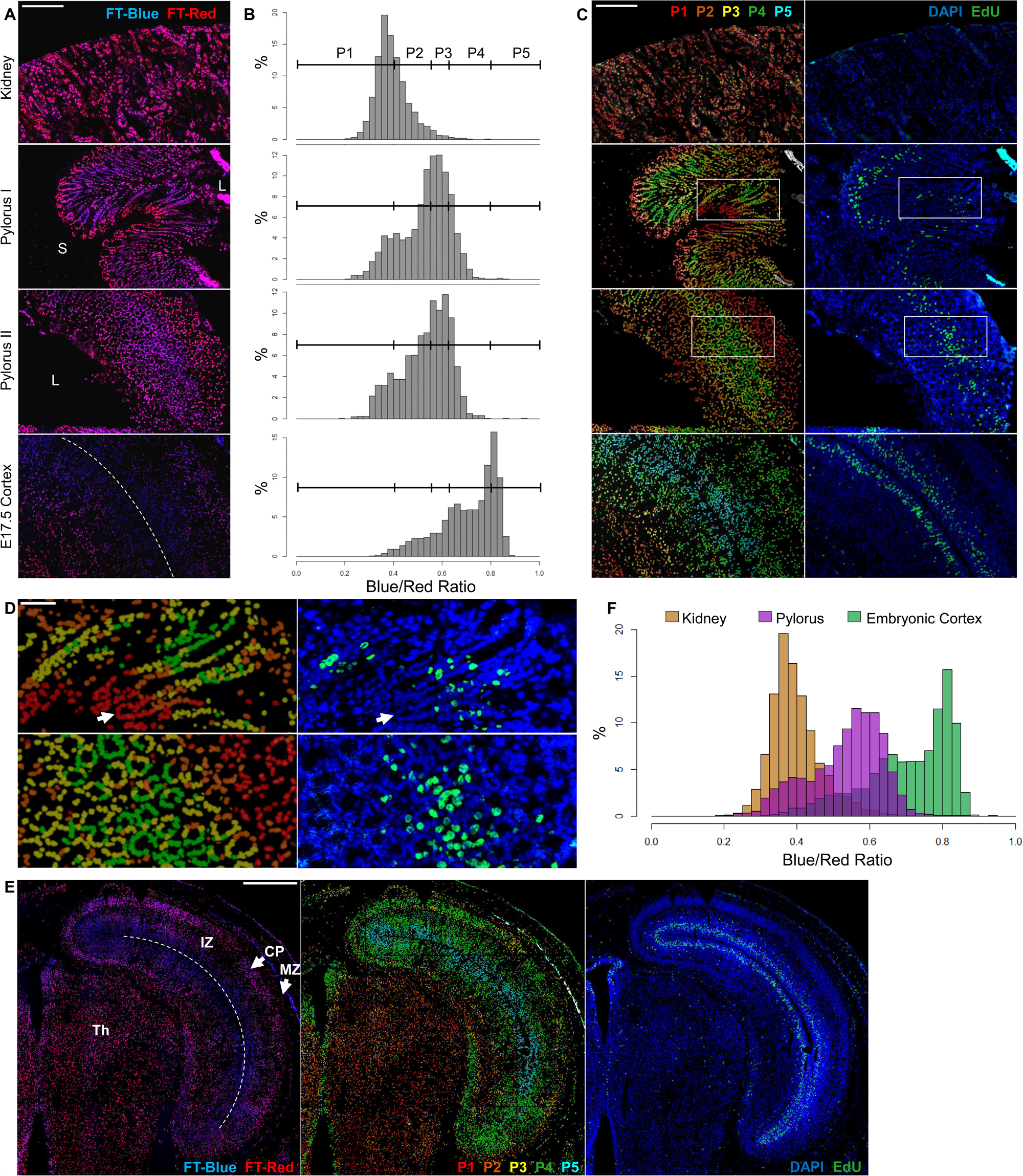
H2B-FT blue/red ratio is consistent with relative turnover rates in solid tissue sections. **A.** 5µm frozen sections from kidney and pylorus in two orientations of an adult iH2B-FT reporter mouse were scanned by microscopy to capture blue and red fluorescence. Gastric glands are shown in elongated orientation (pylorus I) and oblique/cross-sectional orientation (pylorus II). Bottom, the neocortex of a representative E17.5 iH2B-FT mouse embryo. L= pyloric lumen; S=submucosa. Dotted line in bottom left image follows the line of a cerebral lateral ventricle. **B.** The blue/red ratio for each image shown in (A) was quantified and plotted as a histogram, from which five populations of increasing blue/red ratio (P1-P5) were determined. **C.** Left, a heatmap corresponding to populations P1-P5 as shown in (B) was applied onto the original images (A) to indicate the relative locations occupied by cells of different blue/red ratios. Right, the same regions as imaged after fixation and DAPI/EdU labeling. EdU was injected 35 minutes before harvesting mice/embryos. **D.** Magnified detail of white boxed pyloric regions shown in (C). White arrow indicates the base of a gastric gland. **E.** Tiled images showing a zoomed-out view of an E17.5 embryonic brain hemisphere. Left, H2B-FT blue/red merged image. Middle, heatmap of blue/red ratio binned from histograms as in (B). Right, same region imaged after DAPI/EdU labeling. Dotted line indicates cerebral lateral ventricle; IZ=intermediate zone, CP=cortical plate, MZ=marginal zone, Th=thalamus. **F.** Blue/red ratio histograms from all three tissues overlaid on the same plot. “Pylorus” histogram includes combined data from both images of the pylorus (middle images, A). All images contained within a single panel are shown at the same scale. Scale bar in (A,C) = 200µm. Scale bar in (D) = 80µm. Scale bar in (E) = 500µm.

To test the utility of H2B-FT in embryonic development, we examined the developing cortex, another tissue with well-recognized zones of proliferation and differentiation (Sun and Hevner, 2014; Uzquiano et al., 2018). Ventricular radial glia undergo constant asymmetric cell divisions during corticogenesis, while their differentiated progeny undergo a limited number of fate-specifying cell divisions as they migrate away from the ventricular zone (Uzquiano et al., 2018). In frozen sections from E17.5 iH2B-FT embryos, cells in the ventricular zone had the highest blue/red ratio, followed by cells of the subventricular zone (Figure 6A-C, bottom). Cells farther from the ventricle showed a decreased blue/red ratio, particularly underneath the ventricle toward the interior of the brain. When tiled images are stitched together spanning a large tissue area, the H2B-FT color profile reveals distinct layers in the embryonic cortex (Figure 6E). The proliferative behavior revealed by H2B-FT confirms known developmental timelines (Chen et al., 2017) and again is consistent with the pattern of EdU staining (Figure 6C, bottom). Even for cells that exit cell cycle upon differentiation, the gradually decreasing blue/red ratio should – up to a point - report relative time elapsed since their last division. Past this point, cells plateau in their ability to become redder over time, reaching the resolution limit of the H2B-FT reporter. This is exemplified in the homogeneously red-shifted thalamus (Figure 6E), where new neuron production peaks at E14 but trails off by E17 (Chen et al., 2017). Therefore, H2B-FT nuclei quantified *in situ* readily reveals the distribution of relative cell cycle lengths in a given tissue (Figure 6F), similar to what we have shown using flow cytometry. The concordance of the H2B-FT color profile with expected proliferation patterns demonstrates that the reporter on its own can convey cell cycle length. Taken together, the H2B-FT reporter can reveal cell cycle speed *in situ* within the native tissue context.

### H2B-FT blue/red ratio provides an estimate of cell cycle length *in vivo*

Since the H2B-FT blue/red ratio qualitatively agreed with known proliferative behaviors across all biological settings tested, we sought to establish a workflow for quantifying cell cycle length from the reporter readout. Equation S2 describes a mathematical relationship between the steady-state blue/red ratio, 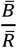, and the cell cycle length, *τ*_*D*_:

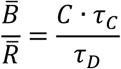

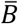 and 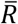 are readily quantified in a fluorescence assay. Therefore, the equation suggests that the cell cycle length *τ*_*D*_ can be calculated as long as the constant *C* is known, since the color conversion time *τ*_*C*_ remains the same for a given FT variant (Figure S1A). The equation also states that C can be derived if *τ*_*D*_ can be experimentally determined. Therefore, we set out to characterize both cell cycle length distribution (Figure 7A) and blue/red ratio distribution (Figure 7B) in the same cells using a combined live imaging + flow cytometry workflow. The details of this workflow are described in the Mathematical Appendix, but in summary, we used our knowledge of ground-truth cell cycle lengths from image-based tracking to solve for *C* · *τ*_*C*_.

**Figure 7.**
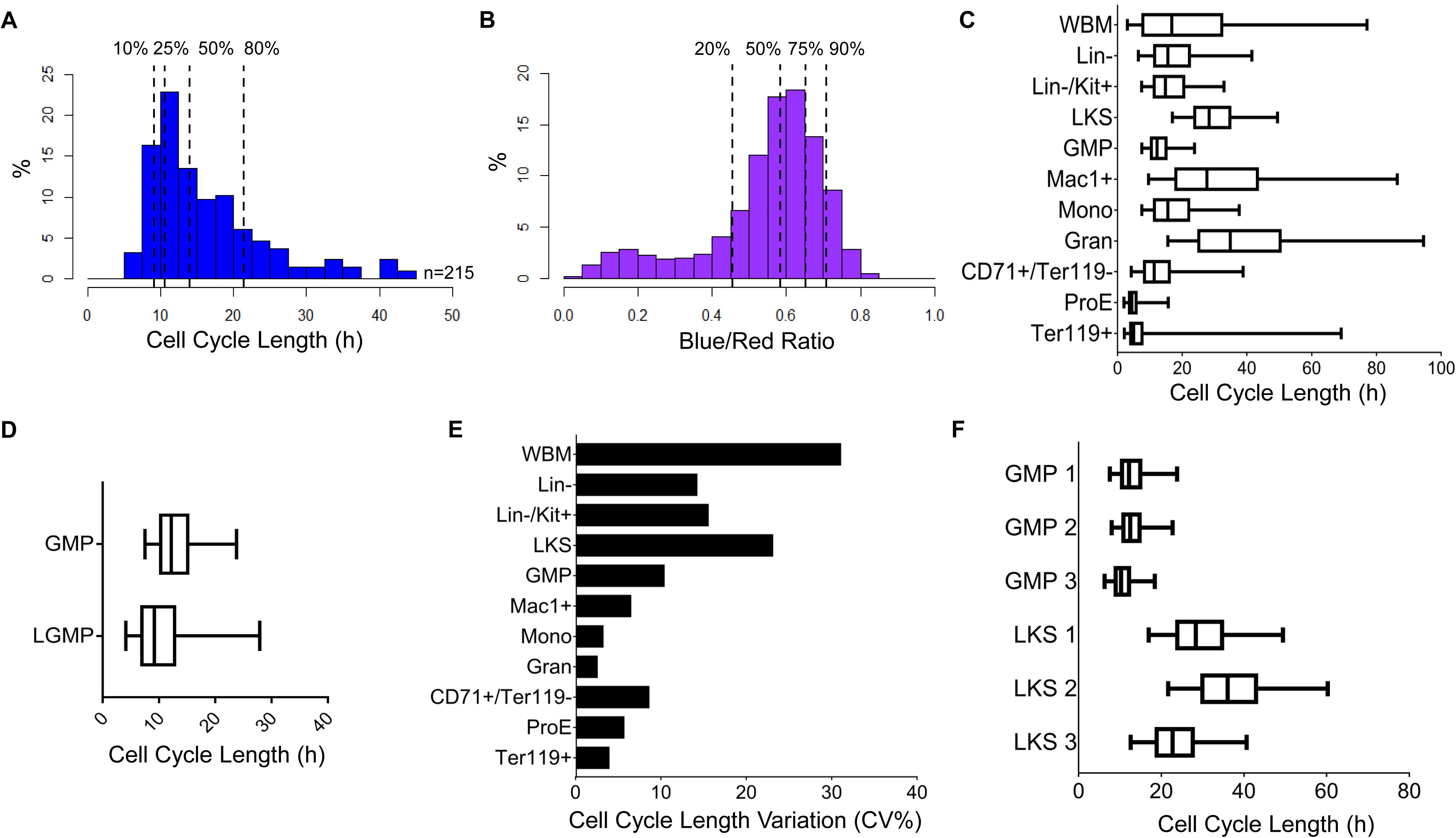
Estimating *in vivo* cell cycle length from H2B-FT blue/red ratio. **A.** Cell cycle length distribution of cultured GMPs. Individual cell cycle length was determined by time-lapse microscopy. Dotted lines show the indicated percentiles. **B.** H2B-FT blue/red ratio distribution of the same GMPs as analyzed by flow cytometry after 24 hours culture. **C.** Calculated cell cycle length distributions of designated bone marrow populations from a representative H2B-FT mouse. **D.** Cell cycle length distribution of normal GMPs and L-GMPs from a representative mouse induced to express MLL-ENL. **E.** Median cell cycle length variation among three H2B-FT mice, shown by percent coefficient of variation (CV%). **F.** Cell cycle length distributions of LKS and GMP populations from the three mice. Boxes in panels (E), (F), and (H) represent the median and interquartile range of each group; whiskers represent 5th-95th percentile.

Having determined the value of *C* · *τ*_*C*_ using cells whose cell cycle length was determined by imaging (Figure 7A-B), we then plugged this constant back into the equation above to obtain cell cycle lengths for various cell types *in vivo*, using fresh bone marrow. Specifically, we assessed the H2B-FT blue/red ratio in mature monocytes and granulocytes (Lagasse and Weissman, 1996) as well as immunophenotypically defined stages of erythropoiesis (Koulnis et al., 2011) (Figure S5F-G, gating strategy). Overall, granulocytes (Gran) were estimated to have an average cell cycle length of >30 hours/cycle. The erythroid progenitors (ProE) appeared to have the shortest cell cycle, averaging at ∼4.3 hours/cycle (Figure 7C). This result was particularly compelling, since it agrees with fetal liver ProE cell cycle length previously determined using sequential thymidine labeling (Hwang et al., 2017). L-GMPs from iMLL-ENL mice had a wider range of cell cycle speeds than normal GMPs, with the median shifted toward a shortened cell cycle length of ∼9 hours/cycle (Figure 7D). For most examined populations, cell cycle length distributions were similar from animal to animal, and the variability generally decreased as the population became more immunophenotypically defined (Figure 7E). High variability was seen in whole bone marrow, which was expected for a heterogeneous tissue (Figure 7E). Strikingly, the LKS compartment displayed high variability across different individuals (Figure 7E), contrasting the close similarity of GMPs from the same cohort. For example, while the GMPs had a median cell cycle length of ∼11.6 ± 1.2 hours/cycle in a cohort of three mice, the median cell cycle length of the corresponding LKS compartment was 22.7, 28.3, and 36.1 hours/cycle (Figure 7F). We note that the median cell cycle length of the LKS compartment is determined by the division rate of the predominant population, which are the more proliferative multipotent progenitors (Passegue et al., 2005). The stem cell-enriched LKS compartment is known to be internally heterogeneous (Copley et al., 2012; Haas et al., 2018; McKenzie et al., 2006). This heterogeneity fluctuates diurnally (Méndez-Ferrer et al., 2009); remodels in response to injury (Haas et al., 2018); and evolves with aging (Beerman et al., 2010; Kowalczyk et al., 2015). Our reporter reveals an unexpected level of heterogeneity across individuals under homeostatic conditions. We anticipate that the H2B-FT mouse offers a unique tool for unveiling how stem and progenitor cell heterogeneity responds to demands placed on the tissue during regeneration, aging, and disease.

## Discussion

Taking advantage of the distinct life spans of a reporter molecule when it exists in two fluorescent states, we describe a novel method for resolving live cell cycle length, *in vitro* as well as *in vivo*. We have shown that cell cycle length can be determined from snapshot measurements by imaging or flow cytometry: the ratio between the blue/red fluorescence intensities faithfully reflects the proliferative state of all cell types tested. Determining cell cycle length *in vivo* has previously been challenging because direct imaging for most tissues is prohibitive. Estimating proliferativeness in deep tissues is largely limited to analyzing relative frequency of cells positive for markers of specific cell cycle phases, mostly S- or M-phases. Indeed, we show that proliferative regions can be grossly demarcated by their high density of EdU+ cells, and they co-localize *in situ* with regions of high blue/red ratio (Figure 6C-E). Although this side-by-side comparison validates the H2B-FT reporter, it also reveals the limitations of the traditional labeling approach. As exemplified in Figure 6C-D, many blue (fast-cycling) cells are EdU-negative because the label only marks cells that chanced to be in S-phase during the pulse. The H2B-FT color profile not only identifies all of the fast-cycling cells in and around the EdU+ population, but additionally reveals gradations in cell cycle length outside the hotspot of EdU labeling. More generally, interpretation of pulse-label data depends on the assumption that S-phase duration is less variable than the gap phases. In cases where cell cycle accelerates by S-phase shortening (Hwang et al., 2017), a fast cell cycle could - counterintuitively - present as decreased frequency of label-positive cells. Conversely, a slowly cycling population arresting at S-phase could present with increased label-positive cells. These shortcomings are partly remedied by sequential labeling with two thymidine analogs (Bokhari and Raza, 1992; Monette et al., 1968). However, the accuracy of the double labeling method depends on population homogeneity because it calculates average cell cycle length of the entire population. In contrast, as H2B-FT reports the blue/red ratio for each cell, it unlocks the intercellular heterogeneity invisible in the sequential thymidine labeling method. The H2B-FT reporter is a first-of-its-kind tool, revealing the distribution of cell cycle lengths inside deep tissues.

The H2B-FT reporter resolves *in vivo* cell cycle speed in a single snapshot, a capability beyond what live cell cycle phase reporters (Bajar et al., 2016; Sakaue-Sawano et al., 2008) can accomplish. Besides revealing relative proliferation rate, the blue/red fluorescence intensity can produce quantitative estimates of cell cycle length after a calibration step to determine the constant *C* · *τ*_*C*_ (Figure 7A-B, Figure S7). The cell cycle lengths determined in this manner agrees well with those determined by double thymine label for the erythroid progenitors (Hwang et al., 2017). We note that the relationship between blue/red ratio and cell cycle length is nonlinear (Figure 1C). Experimental data from MEFs and cultured GMPs, where both cell cycle length and blue/red ratio were measured, supported the model and showed the resolution plateauing beyond approximately 30 hours/cycle (Figure 3F and Figure S7J). Although flow cytometry could help extend this dynamic range owning to its superior sensitivity, this H2B-FT reporter is expected to be more suitable for resolving faster-cycling cells. For improved resolution at longer cell cycle length, future versions of the reporter could include using the slow FT variant (Figure S1A, Figure S3C-D). Additionally, synchronizing cells from their most recent mitosis – for example, with the help of a cell cycle phase reporter - could help correct for the predicted “breathing” of the blue/red ratio across the duration of a single cell cycle (Figure S1).

We anticipate wide applicability of the H2B-FT as cellular heterogeneity is better appreciated with the advent of single cell genomics (Battich et al., 2015), particularly when cell cycle is known to be a major driver of this heterogeneity (Olsson et al., 2016). With this reporter targeted into the mouse genome, even cell types as rare as hematopoietic stem cells can be reliably analyzed (Figure 5,7). As the blue/red fluorescence is compatible with additional fluorescent markers, this allele is suitable to be crossed with GFP-expressing reporters to further refine cells of interest. Unlike assays based on label dilution (Falkowska-Hansen et al., 2010; Lyons et al., 2001; Tumbar et al., 2004; Wilson et al., 2008), the readout from the genetically encoded H2B-FT does not decay over cellular generations. We have also demonstrated that the H2B-FT can be conveniently expressed, e.g. by a viral vector (Figure 2-5), to enable the assessment of cell cycle rate in other cellular models, including human cells. The ability to physically separate live cells free of additional labels or perturbations provides new opportunities for functional and mechanistic studies on how cell cycle dynamics regulate the specification, maintenance, and/or reprogramming of cellular identity. As one example, our previous work demonstrated the extraordinary response of naturally fast-dividing GMPs (∼8 hours per cell cycle) to pluripotency induction (Guo et al., 2014) and to malignancy initiation (Chen et al., 2018). Sorting the fastest-cycling GMPs using the H2B-FT reporter would provide an unprecedented opportunity to look into a molecular state from which pluripotency and malignancy could emerge essentially unopposed. H2B-FT should also be helpful in elucidating mechanisms of cell fate control during development. For example, it is widely thought that the balance between cell proliferation and differentiation is critical in the development of the neocortex, and that dysregulation of either one of these processes carries severe neurological consequences (Ernst, 2016). A sensitive, continuous monitor of cell cycle speed would therefore be a valuable tool in the study of neurodevelopment and its associated disorders. Finally, the method described here should also be applicable to questions of more immediate clinical relevance, such as the functional differences between fast- and slow-dividing cancer cells in terms of metastatic potential, therapy response, and immune evasion. When expressed specifically in the host, the H2B-FT could also reveal potential proliferation heterogeneity in the complex tumor niche and immune surveillance system (Madar et al., 2013; Solito et al., 2014). Overall, the contribution by the speed of cell cycle to diverse biology is expected to unravel with the H2B-FT.

Beyond its technical utility, the mathematical principles underlying the H2B-FT reporter have important biological implications. The experimental H2B-FT data depict a generalized relationship between molecular half-life and intracellular concentration. These findings constitute direct evidence for a mechanism by which cells can selectively alter their molecular contents by modulating cell cycle dynamics. Simplistically, intracellular molecules can be placed in two categories: those that exist long enough such that their intracellular concentration is dependent on cell cycle length; and shorter-lived molecules which are less sensitive to dilution by cell division. Since these principles do not discriminate against any specific types of molecule, they suggest a fundamental mechanism by which cell cycle speed might be tuned to optimize the concentration of key molecular regulators, including proteins, their various post-translationally modified derivatives, RNA species, and others. This aspect of regulation could echo the direct regulation on the half-life of proteins and RNAs, since the effective concentration is determined not only by the turnover rate, but also by the cell cycle length.

## Mathematical Appendix

This appendix describes the details of cell cycle length quantification from the blue/red ratio readout of the H2B-FT reporter.

### Determining the equation constant

Equation S2B describes a mathematical relationship between the steady-state blue/red ratio, 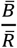, and the cell cycle length, *τ*_*D*_:

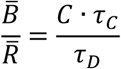

where *τ*_*C*_ is the time required for the molecular conversion of blue to red, and *C* is a normalization constant. Because 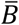 and 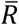 are readily quantified in a fluorescence assay such as FACS, the equation suggests that the cell cycle length *τ*_*D*_ can be calculated if the constant *C* can be experimentally determined, as the color conversion time *τ*_*C*_ remains the same for a given FT variant (Figure S1A). With the 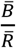 value normalized to the total cellular FT content, 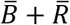, the modified Equation S2B then becomes:

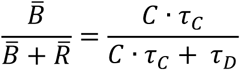

We found that *C* depended on the fluorescence detection parameters: exposure time in the case of fluorescence microscopy, and laser voltage for flow cytometry (Figure S7A-E). To determine the value of *C*, we directly measured the cell cycle lengths of iH2B-FT GMPs during a 24-hour culture period by live-cell imaging and single-cell tracking (Figure 7A). After 24 hours, the imaged iH2B-FT GMPs were analyzed by FACS to determine their blue/red profile (Figure 7B). Thus, the blue/red ratio distribution for these 24-hour cultured cells could be matched to their ground-truth cell cycle speed distribution. This provided paired coordinate values for 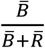 and cell cycle length *τ*_*D*_ such that the value of the combined constant *C* · *τ*_*C*_ could be calculated from Equation S2B (Figure S7F-G). Separate calculations were performed using coordinates representing the 10^th^, 25^th^, 50^th^ and 80^th^ percentile of cell cycle length, assumed to correspond to the 90^th^, 75^th^, 50^th^ and 20^th^ percentile of the bluest cells, respectively (Figure S7G). The resulting C · τC values were then plugged back into Equation S2B to produce curves which we evaluated for their similarity to each other, and to the experimental microscopy data (Figure S7H). Based on the plot in Figure S7H, a *C* · *τ*_*C*_ value representing the average of predictions 2 and 3 was selected for estimating *in vivo* cell cycle lengths.

### Consideration of cell-division-independent H2B-FT-red decay

As H2B-FT-red does exhibit decay independent of cell division in MEFs and HeLa cells (Figure S2), we considered a modified version of Equation S2 in which a rate constant for active degradation, τ_R_, was incorporated into the red removal rate, δ (Figure S7I):

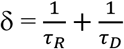

However, as the exact red degradation parameter in hematopoietic cells *in vivo* is unknown, we examined the relationship between blue/red ratio and cell cycle length by modeling a wide range of red decay rates (Figure S7J-K). These exercises showed that for cell cycle lengths shorter than 30 hours, the red degradation rate was irrelevant; there was little difference in cell cycle lengths whether H2B-FT-red was assumed to be shorter (35 hours) or longer (135 hours), or whether it was simply ignored 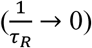, when δ becomes simply inversely proportional to τ_D_ as in Equation S2B. At longer cell cycle lengths (>30h), ignoring red decay resulted in the shortest possible prediction of cell cycle length for a given blue/red value, while incorporating red decay adjusted these predictions upward in a manner inversely correlated with half-life (Figure S7J-K). To visualize these different scenarios in our actual bone marrow data, we proceeded to generate two estimates of cell cycle length with different red decay parameters: either by ignoring cell division-independent decay or by setting the red fluorescence half-life to 85h (Figure S7L). Although absolute cell cycle length for slow-dividing cells remains more uncertain than in fast-cycling cells, the relative differences between cell types are still maintained in any version of the equation. Therefore, in order to compare cell cycle length distributions across many hematopoietic cell types and different individual mice (Figure 7C-F), we chose the simplest version of the equation (red decay ignored) for reporting results, while acknowledging that slow cell cycle lengths (>30h) are likely to be underestimated using this assumption.

With these considerations and having determined the value of *C* · *τ*_*C*_, we proceeded to estimate the cell cycle length distributions for various hematopoietic cell types (Figure 7C).

## Methods

### Cloning and reporter cell line generation

pFast-FT-N1, pMedium-FT-N1, and pSlow-FT-N1 were obtained from Addgene (31910, 31911, and 31912 respectively). Each FT coding sequence (∼711 bp each) was cloned into a Dox-inducible lentiviral backbone and a constitutive retroviral backbone. The inducible lentiviral plasmid (pFU-TetO-Gateway-PGK-Puro) was constructed previously(Guo et al., 2012) by inserting a Gateway cassette (Thermo Fisher Scientific/Invitrogen), a PGK promoter, and a puromycin resistance gene into the pFU-tetO-Klf4 vector(Stadtfeld et al., 2008) through blunted EcoRI sites. Each FT insert was then cloned into this destination vector through Gateway recombination. For constitutive expression, the FT sequences were inserted into a pSCMV(Guo et al., 2012) retroviral backbone using HindIII and XhoI restriction sites. The H2B-FT fusion transgenes were constructed by overlap extension PCR using the three FT plasmids as well as the human H2B.J coding sequence from PGK-H2B-mCherry (Addgene # 21217) as templates, and cloned into the pFU-TetO-Gateway-PGK-Puro and pSCMV expression plasmids. H2B-mCherry and H2B-BFP (BFP template from Addgene #52115) inserts were similarly cloned to serve as single-color controls for the color-changing H2B-FT.

The MSCV-IRES-GFP (“empty vector”) retroviral expression plasmid was previously described(Lu et al., 2008). The LZRS-c-Myc-IRES-GFP retroviral expression plasmid was a gift from Sebastian Nijman.

Viral vectors were transfected in 293T cells using Fugene® 6 transfection reagent (Promega). Viruses harvested from the supernatant were used to transduce BaF3 and HeLa cells. Successfully transduced cells were selected by FACS.

*HPRT::*iH2B-FT knock-in mouse embryonic stem cells (mESCs) were generated using inducible cassette exchange (ICE) to target the TetO-CMV-H2B-FT transgene to the *HPRT* locus by cre-recombination in the A2lox.cre mESC cell line (Iacovino et al., 2011). Briefly, H2B-FT inserts (all three kinetic variants, Subach et al. and Figure S1A), as well as H2B-BFP and H2B-mCherry were cloned into P2lox targeting plasmids (Iacovino et al., 2011) using HindIII and NotI restriction sites. The targeting plasmids were electroporated into Dox-activated A2lox.cre mESCs. After one day of recovery on neomycin-resistant feeder MEFs (Millipore Sigma), successfully recombined clones were selected by supplementing the mESC culture medium with 300µg/mL geneticin (G418). After 6 days of selection, healthy surviving colonies were hand-picked under an inverted microscope and replated onto WT irradiated feeder MEFs. These “Passage 0” cultures were subsequently split for cryopreservation as well as continued culture and characterization of reporter activity by fluorescence microscopy and flow cytometry.

mESCs were made to constitutively express H2B-FT under an EF-1α promoter using the Sleeping Beauty transposon system (Hudecek et al., 2017). H2B-FT inserts (all three variants) were cloned into a modified empty transposon (pT3) backbone. A GFP sequence in the original backbone, pT3-Neo-EF1a-GFP (Addgene # 69134), was replaced with a multiple cloning site (MCS) synthesized by IDT using MluI and NotI restriction cloning. The H2B-FT sequence was then inserted into the MCS using PmeI and NotI restriction sites. pT3-EF1a-H2B-FT vectors were transfected into WT mESCs using Lipofectamine™ 2000 reagent (Invitrogen) along with a separate vector, pCMV(CAT)T7-SB100 (Addgene #34879), encoding the Sleeping Beauty transposase. Successfully transfected mESCs were enriched by FACS.

Primer sequences used for cloning are provided in Table S1. Plasmid maps are available upon request.

### Cell culture and mESC differentiation

MEFs, HeLa, and 293T cells were cultured in a standard growth medium (“MEF medium”) consisting of DMEM basal medium (Gibco) with 10% heat-inactivated fetal bovine serum (FBS) (Gibco) and 1% penicillin/streptomycin/L-Glutamine supplement (ThermoFisher). For MEF derivation from E13.5 embryos, this medium was additionally supplemented with 1% Non-essential amino acid (NEAA) mixture (ThermoFisher) for the first passage *in vitro*. 293T cells used for viral transfection were cultured in the standard growth medium additionally supplemented with 1% sodium pyruvate (ThermoFisher). Occasionally, phenol-red-free Medium 199 was used in place of DMEM for live fluorescence microscopy experiments with H2B-FT HeLa cells (details in main text). Mouse embryonic stem cells (mESCs) were cultured either on irradiated feeder MEFs or on plates coated with 0.1% gelatin in DMEM supplemented with 15% ESC-qualified FBS (Millipore), 1% penicillin/streptomycin/L-Glutamine, 1% NEAA, 1000U/ml LIF (Millipore), and 0.8µl/100mL β-mercaptoethanol. BaF3 cells were cultured in an RPMI-based growth medium consisting of 10% heat-inactivated FBS, 1% penicillin/streptomycin/L-Glutamine, and 270pg/mL IL-3 (Peprotech). Primary GMPs were cultured in complete X-vivo medium (Lonza) supplemented with 10% BSA (Stemcell Technologies), 1% penicillin/streptomycin/L-Glutamine, 0.14µl/mL β-mercaptoethanol, and 100 ng/ml mSCF, 50 ng/ml mIL3, 50 ng/ml Flt3L, and 50 ng/ml mTPO (all from PeproTech).

*In vitro*, doxycycline (Sigma) was added to cell culture medium at 2µg/mL for inducible promoter activation. To induce iH2B-FT and/or iMLL-ENL expression *in vivo*, mice were fed drinking water containing 1g/L Dox supplemented with 10g/L sucrose.

For the differentiation assay, mESCs maintained on Feeder MEFs were transferred to feeder-free conditions (0.1% gelatin) for 2 passages to potentiate exit from pluripotency. The pluripotency of control cells was maintained with mESC culture medium, while the differentiation condition entailed switching the cells into standard MEF medium supplemented with 2µM retinoic acid (RA) (Sigma).

For the colony recovery assay, sorted cells were plated onto feeder MEFs (n = 4 replicate wells) at a standardized seeding density and fed with mESC medium. After 6 days, the cultures were fixed and alkaline phosphatase (AP) staining was done using the Stemgent APII kit in accordance with the protocol provided by the manufacturer. Colonies staining positive for AP activity were counted manually (RA treated group) or using an automated image processing workflow (mESCs).

Total mRNA was extracted with TRIzol® Reagent (Ambion) and reverse transcribed into cDNA using SuperScriptIII™ First-Strand Synthesis SuperMix (Invitrogen) according to the product manual. For quantitative real-time PCR, cDNA and gene-specific primers were mixed with iQ™ SYBR®Green Supermix (Bio-Rad) and carried out using a Bio-Rad CFX384™ Real-Time PCR System. Gene expression levels were normalized to GAPDH level in the same sample. qPCR primer sequences are provided in Table S1.

### Mice

All mouse work was approved by the Institutional Animal Care and Use Committee (IACUC) of Yale University. All research animals were housed and maintained in facilities of Yale Animal Resource Center (YARC).

*HPRT*::iH2B-FT-Medium and *HPRT*::iH2B-FT-Slow chimeric mice were generated from passage 0 A2lox.cre H2B-FT targeted mouse embryonic stem cells by the Yale Genome Editing Center via blastocyst injection and implantation into C57/Bl6 females. High-degree chimeric male offspring were selected by coat color and the iH2B-FT allele was subsequently backcrossed onto a C57/Bl6 background. For all H2B-FT blue/red analysis of primary-harvested cells, H2B-FT expression was induced *in vivo* for at least one week by feeding Dox drinking water. Tissues from iH2B-FT mice were harvested and analyzed when the mice were 8-12 weeks of age. The iH2B-FT x iMLL-ENL mice were harvested at 5 weeks old. All cohorts reported here involved male mice.

WT feeder MEFs for pluripotent stem cell culture were derived from C57/Bl6 (Jackson Lab) E13.5 mouse embryos and mitotically inactivated by 80 Gy γ-irradiation. H2B-FT-Medium MEFs were derived from *HPRT*::iH2B-FT-Medium E13.5 mouse embryos.

### HSPC transplantation

FACS-purified bone-marrow-derived LKS cells from donor mice were transduced overnight *in vitro* with TetO-H2B-FT lentivirus and injected through the tail vein at >10,000 cells/mouse into 9-week-old γ-irradiated recipient mice (9 Gy) along with 500,000 WBM support cells. Alternatively, donor mice were injected intraperitoneally with 150µg/g body weight 5-fluorouracil (5-FU) 4 days before harvesting, and the HSPC-enriched WBM was virally transduced and then transplanted at a ratio of 1 donor per 2 recipients. Transplanted mice were given a one-time intraperitoneal injection of 200µg Dox and thereafter continuously maintained on Dox drinking water. After four weeks, peripheral blood was collected and analyzed by flow cytometry to evaluate H2B-FT expression coming from the engrafted virally transduced cells. Bone marrow HSPCs were harvested between 8-16 weeks post-transplantation and stained with fluorescent antibodies marking Lineage, Kit, Sca-1, CD34, and CD16/32. Cellular fluorescence was recorded on an LSRII flow cytometer (BD) or a FACSAria™ cell sorter (BD) using FACSDiva™ software (BD), and the data were subsequently analyzed using FlowJo software (FlowJo, LLC).

A complete list of antibodies used in this study is provided in Table S2.

### EdU/DAPI labeling

Cultured cells were treated with growth medium containing 10µM EdU for 15 minutes. EdU was rinsed away with PBS and cells were trypsinized into a single-cell suspension for immediate fixation or H2B-FT blue/red FACS-sorting followed by fixation. For *in vivo* labeling of adult tissues, mice were pulsed with EdU at 50µg/g body weight via intraperitoneal injection either 2 hours (Figure 5F) or 35 minutes (Figure 6) before harvesting. Pregnant dams were given EdU intravenously at 50µg/g body weight 35 minutes before harvesting embryos (Figure 6, E17.5 cortex).

H2B-FT-blue and red signal was captured (by FACS or microscopy) in unfixed cells/tissue prior to downstream steps, as exposure to fixative prematurely converts blue molecules into red.

Labeling in suspension: Bone-marrow-derived HSPCs plus Ter119+ cells were enriched via streptavidin-conjugated magnetic microbead separation (Miltenyi Biotec, Product #130-048-101) of lineage+ cells (lineage cocktail included biotin-conjugated CD3e, CD4, CD8, CD11b, Gr1, and B220), stained with GMP markers or CD71/Ter119, and sorted by FACS on a FACSAria™ (BD) directly into 70% ethanol. These fixed single-cell suspensions of EdU-treated cells were stored in 70% ethanol at −20C for >24h, and then rinsed in PBS. Cells were permeablized in 0.2% Triton X-100 in PBS at room temperature for 15 minutes and then fluorescently labeled by Click chemistry using a Click-IT EdU-488 kit (ThermoFisher Scientific, Product #C10337) for 30 minutes at room temperature in an AF488-azide EdU labeling cocktail prepared according to the product manual. Cells were rinsed in PBS and incubated for 10 minutes at room temperature in 1µg/mL DAPI (ThermoFisher Scientific #D1306) diluted in PBS. Cells were rinsed once more with PBS and resuspended in a PBS buffer containing 1% BSA, then analyzed by flow cytometry on a BD LSRII.

Labeling on slides: Intact organs (kidney, stomach) and whole embryos were embedded in cryomolds (Sakura) filled with OCT compound (Sakura), and flash-frozen in a 2-methylbutane/liquid nitrogen bath. 5µm sections were cut at −25C using a CM3050 S cryostat (Leica) and mounted onto SuperFrost® slides (Fisher Scientific). Frozen sections were scanned by fluorescence microscopy (details below) at 20x to capture blue and red H2B-FT signal, and then slides were fixed in 4% paraformaldehyde (Electron Microscopy Sciences) for 12 minutes at room temperature. Samples were washed, permeablized, and stained as described above.

### Microscopy

Live- and fixed-cell microscopy was performed using a Molecular Devices ImageXpress® Micro 4 high-throughput compound inverted epifluorescence microscope equipped with a live imaging environment. For time lapse imaging, cells were plated into a Greiner Bio-One CellStar® 96well plate, sealed with a Breathe-Easy® gas-permeable membrane, and maintained at 37C/5% CO2 for the duration of the experiment. MetaXpress® 6 software was used for all image acquisition and for some image processing and analysis. Most of the image segmentation and image-based measurements were carried out using CellProfiler™ and exported into FCS Express 6 Image Cytometry software (De Novo™ Software) for data analysis. CellProfiler™ pipelines and sample raw images from our experiments are available on request. ImageJ software was used for some image processing.

### Statistical Information

Details pertaining to each statistical analysis are provided in the figure legend accompanying the relevant data figure. All statistical tests used were two-tailed and were carried out using Prism software (GraphPad). All measurements were taken from distinct samples.

### Data and Materials Availability Statement

The data that support the findings of this study are available from the corresponding author upon reasonable request, as are unique biological materials generated by the authors.

## Acknowledgements

This work was funded by a National Institutes of Health award (DP2GM123507-01); the Charles H. Hood Foundation; and Gilead Sciences. A.E.E. was supported in part by the NIH/NIGMS Cellular and Molecular Biology Training Program under award number T32GM007223. The authors would like to thank the Yale Stem Cell Center, the Yale Cooperative Center for Excellence in Hematology, Dr. Diane Krause (for providing feedback and reagents), the Yale Flow Cytometry Core, and the Yale Genome Editing Core.

## Author Contributions

A.E.E. and S.G. conceived and designed experiments. A.E.E., X.C., X.H., A.M.P.M., and S.G. performed experiments. A.E.E., A.M.P.M., and C.Y. analyzed data. X.C., X.H., A.A.H., and J.L. provided samples. H.Y.K. wrote the mathematical equations. A.M.P.M. and J.L. assisted with mathematical modeling. X.C., X.H., A.A.H., J.L., and H.Y.K. assisted A.E.E. and S.G. with critical feedback and data interpretation. A.E.E. and S.G. prepared figures and wrote the manuscript.

## Declaration of Interests

The authors declare no competing interests.

