## Supplement for "Resolving cell cycle speed in one snapshot with a live-cell fluorescent reporter"

**A**

Here, we derive mathematical expressions for differences in the level of a molecule *M* between two cell populations, one with a shorter cell cycle time $\tau_1$, and another with a longer cell cycle time $\tau_2$. The dynamical equations describing the time evolution of this system are given by:

$$\frac{dM_1}{dt} = k_1 - \left(\frac{1}{\tau_1} + \frac{1}{\tau_P}\right) M_1$$

$$\frac{dM_2}{dt} = k_2 - \left(\frac{1}{\tau_2} + \frac{1}{\tau_P}\right) M_2$$

where $M_1$ and $M_2$ give the levels of the molecules in the two cell populations, $k_1$ and $k_2$ give its synthesis rates in the two populations respectively, and $\tau_P$ gives its time constant for degradation, which is assumed to be the same in the two populations. Solving for the steady-state level of the molecular species in the system, we have that:

$$M_1 = k_1 \tau_1 \left(\frac{\sigma_P}{\sigma_P + 1}\right)$$

where $\sigma_P = \tau_P/\tau_1$ is the ratio of the molecular half-life to the cell cycle duration of the first cell population. Similarly,

$$M_2 = k_2 \tau_2 \left(\frac{\sigma_P}{\sigma_P + \sigma_2}\right)$$

where $\sigma_2 = \tau_2/\tau_1 > 1$ is the ratio of the cell cycle duration of the second population to that of the first populations. From these two equations, we find that the ratio of the molecule levels of the slow cycling population to that of the fast-cycling population is given by:

$$\mu = \frac{\kappa\, \sigma_2(\sigma_P + 1)}{(\sigma_P + \sigma_2)}$$

where $\kappa = k_2/k_1$ is the ratio of synthesis rates for the slow versus fast-cycling populations.

From this expression a few features are apparent: 1) when the molecule is extremely unstable compared to the cell cycle time ($\sigma_P \ll 1$), this ratio converges to the ratio of synthesis rates between the two populations $\kappa$. 2) When the molecule half-life is much longer than the longer cell cycle length $\sigma_P \gg \sigma_2$, this ratio is now given by the synthesis rates multiplied by the ratio of the long cell-cycle length to the short cell cycle length $\kappa\sigma_2$. Thus, when synthesis rates of the two populations are the same, relative differences in the abundance of this molecule are simply given by the relative differences in cell cycle lengths between these two populations. When synthesis rates of two populations are different ($\kappa \neq 1$), cell cycle length differences amplify these differences by a constant multiplicative factor. 3) As the half-life of the molecule increases, relative differences in its levels increase, and reach a half-maximum when its half-life is equal to the longer cell cycle length ($\sigma_P = \sigma_2$; see black dot in (b)).

**B**

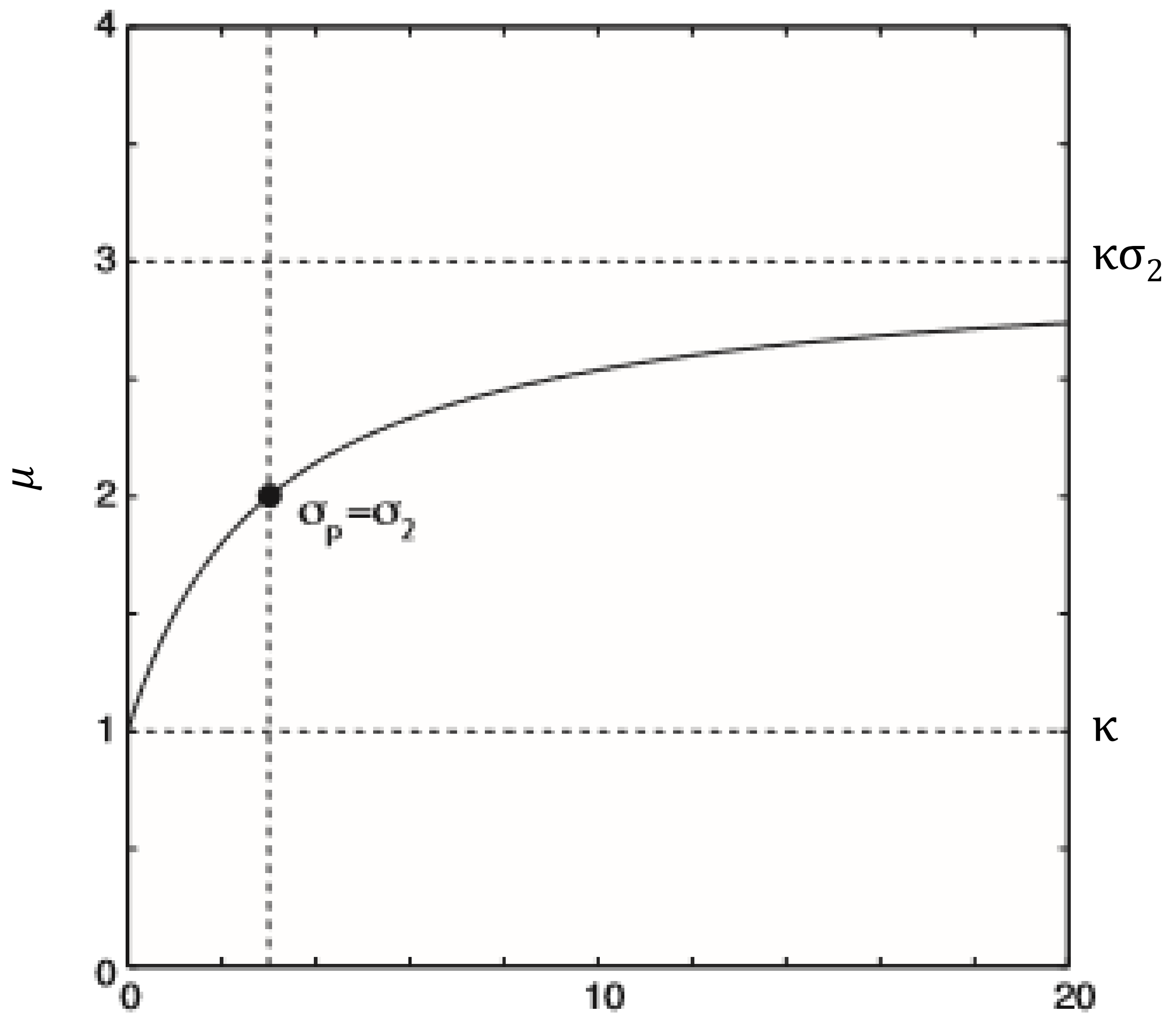


**Equation S1. A molecule's intracellular level depends on the molecule's half-life and cell cycle length. A.** The equation describing the ratio (μ) of levels of a molecule (*M)* in two populations of cells with different cell cycle lengths. $\sigma_2 = \tau_2/\tau_1 > 1$ is the ratio of cell cycle lengths; $\sigma_P = \tau_P/\tau_1$ is the ratio of the molecule's half-life to the shorter cell cycle length; and $\kappa = k_2/k_1$ is the ratio of the molecule's synthesis rates between the two cell populations. **B.** Graph of μ values illustrating the dependence of molecule levels on the half-life and cell cycle length. Parameters: $\kappa = 1$, and $\sigma_2 = 3$. Note that levels of molecules at which this ratio is half-maximal is given by the black dot ($\sigma_P = \sigma_2$).

**A**

The levels of stable fluorescent proteins are strongly influenced by cell division, which results in their removal by dilution. We can exploit this concept to generate a live-cell reporter of cell cycle speed, using a color-changing fluorescent timer protein. The following ordinary differential equations describe the time evolution of such a timer:

$$\frac{dB}{dt} = \alpha - \frac{B}{\tau_C}$$
$$\frac{dR}{dt} = \beta B - \delta R$$

Here *B* is the blue emitting initial molecular species, *R* is the red-emitting molecular species after fluorescence conversion, $\alpha$ is the synthesis rate, $\tau_C$ is the time constant for fluorescence conversion, and $\delta$ is the rate of protein removal. In the most general case, the rate of protein removal is a sum of the rates of proteasomal degradation and cell division; however, when the fluorescent protein is stable, this parameter is simply inversely proportional to the cell cycle time $\tau_D$:

$$\delta = 1/\tau_D$$

Solving for the steady state of the system, we find that:

$$\bar{B} = \tau_C \alpha$$
$$\bar{R} = \alpha/\delta$$

Consequently, we find that the observed steady-state ratio of the two quantities;

$$\frac{\bar{B}}{\bar{R}} = C \cdot \tau_C \left(\frac{1}{\tau_D}\right)$$

where *C* is a normalization constant.

**C**

The rate of protein removal, $\delta$, can be expanded in scope to include active degradation in addition to cell-division-dependent dilution. When both processes are active:

$$\delta = 1/\tau_R + 1/\tau_D$$

where $\tau_D$ is cell division time, and $\tau_R$ is the time constant for active degradation. Solving for the steady state of the system, we find that:

$$\bar{B} = \tau_C \alpha$$
$$\bar{R} = \alpha/\delta$$

Consequently, we find that the observed steady-state ratio of the two quantities;

$$\frac{\bar{B}}{\bar{R}} = C \cdot \tau_C \left(\frac{1}{\tau_D} + \frac{1}{\tau_R}\right)$$

where *C* is a normalization constant.

**D**

$$\frac{\bar{B}}{\bar{R}} = C \cdot \tau_C \left(\frac{1}{\tau_D} + \frac{1}{\tau_R}\right)$$

**B**

$$\frac{\bar{B}}{\bar{R}} = \frac{C \cdot \tau_C}{\tau_D}$$

**Equation S2. Relationship between FT blue/red ratio and cell cycle length. A.** The steady-state levels of the blue and red forms of the FT ($\bar{B}$ and $\bar{R}$, respectively) are the net result of protein synthesis rate ($\alpha$) and turnover rate. Blue turnover ($\beta$) happens upon conversion from blue to red after a period of time ($\tau_C$). Since red molecules are stable, the rate of red removal ($\delta$) can be understood to be inversely proportional to the cell cycle time ($\tau_D$). *C* is a normalization constant. **B.** Taken together, a simplistic estimation of cell cycle length can be determined from the steady-state blue/red ratio or vice-versa. **C-D.** A modified version of the equation incorporates a rate constant, $\tau_R$, for active protein degradation. $\tau_R$ is related to protein half-life via the formula $T_{1/2} = \tau_R \cdot ln2$.

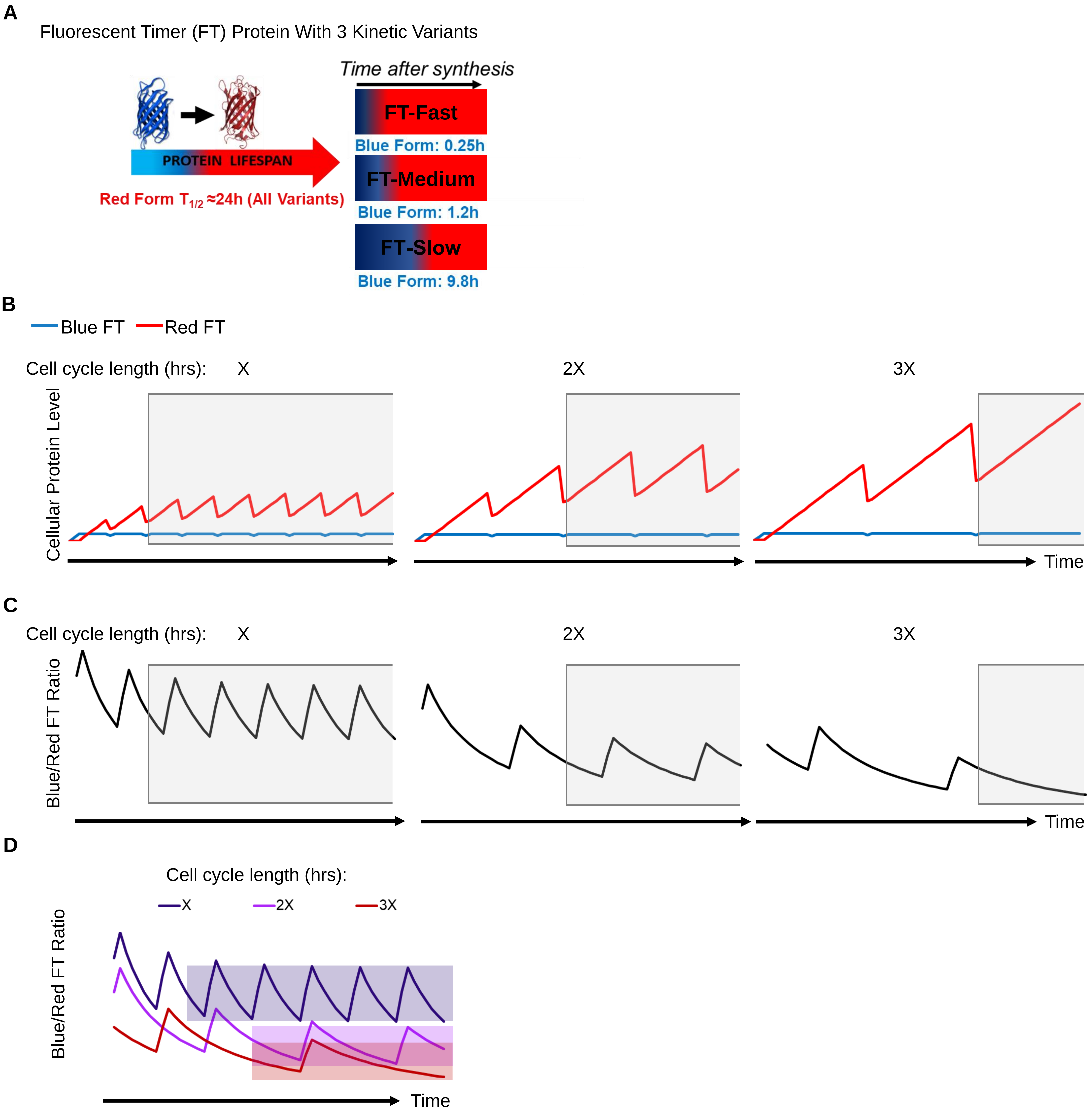


**Figure S1. Modeling the dynamic behavior of FT fluorescence in relation to cell cycle length. A.** The lifespan of the blue (immature) and red (mature) forms of the monomeric fluorescent timer (FT). Three kinetic variants were described by Subach et al. 2009. **B.** Results from mathematical modeling of blue and red FT levels following the onset of gene expression. Gray box shows the FT levels after reaching steady-state. Cells cycling at three distinct speeds are depicted: *X* (left), *2X* (middle) and *3X* (right) hours per cycle. Peaks/valleys represent maximum/minimum FT levels achieved before/after each mitosis. **C.** Blue/red FT ratio of cellular models from (B). **D.** Superimposed plots from (C). Shading indicates the area between the maximum and minimum blue/red ratio for each cell cycle length, showing the anticipated separation of cells with different cycle lengths based on blue/red ratio.

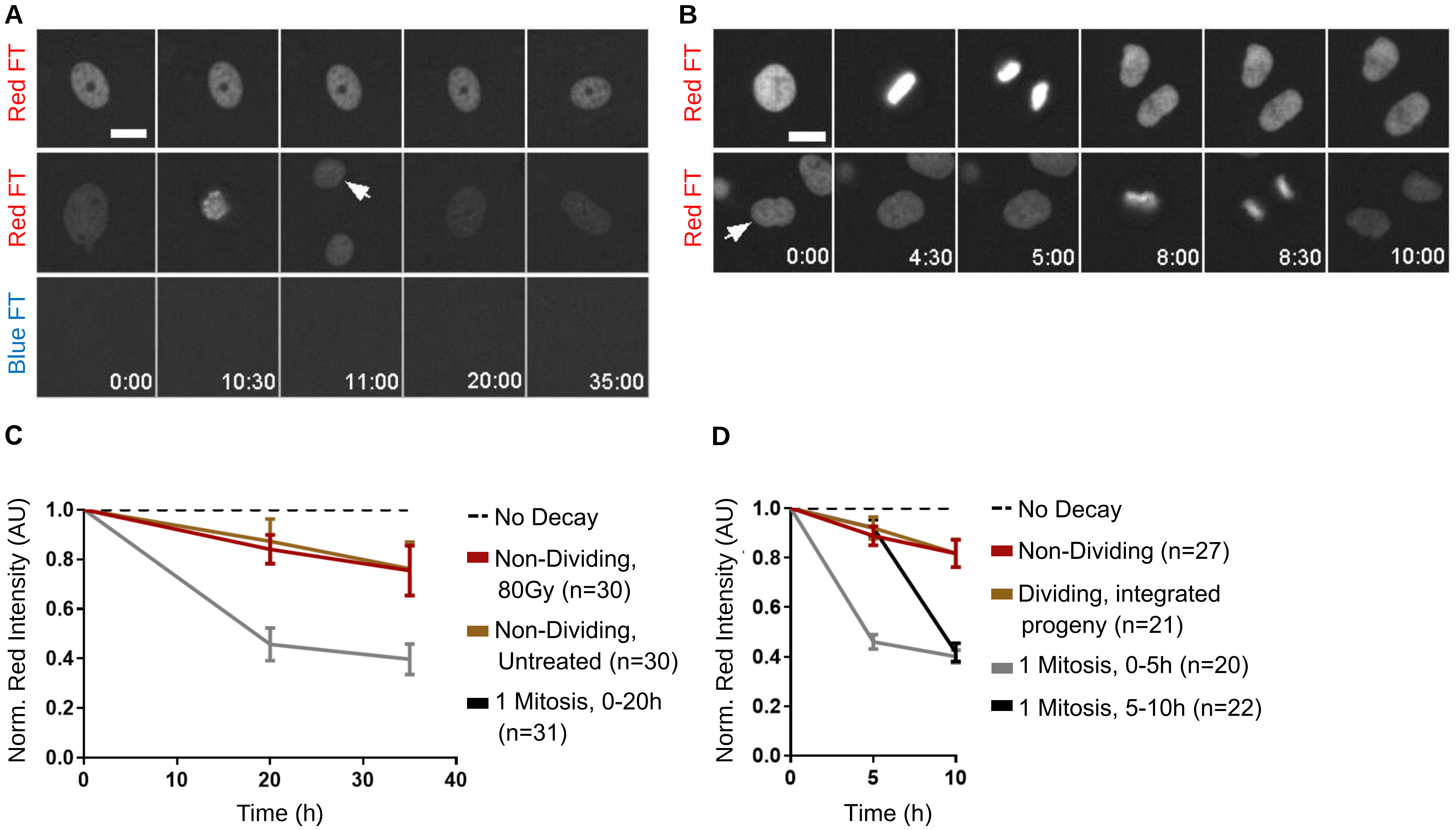


**Figure S2. Characterization of H2B-FT-red decay in cultured cells. A.** Representative time series of primary iH2B-FT MEFs after Dox washout. Cells that never divided (top row) and cells that divided once (middle row) during the imaging period are shown. **B.** Representative time series of Hela cells after Dox washout. Cells that divided early (top row) and cells that divided late (middle row) during the imaging period are shown. **C.** Quantification of the red FT in MEFs over time. Time 0 refers to the point when H2B-FT-blue signal fell below detection limit, due to complete conversion to red. **D.** Quantification of the red FT over time in Hela cells. When tracking the red fluorescence decay, the integrated red intensity measurements of all daughter nuclei were added together. Error bars denote standard deviation; n refers to the number of single cells tracked per condition. Scale bars in (A-B) = 20µm.

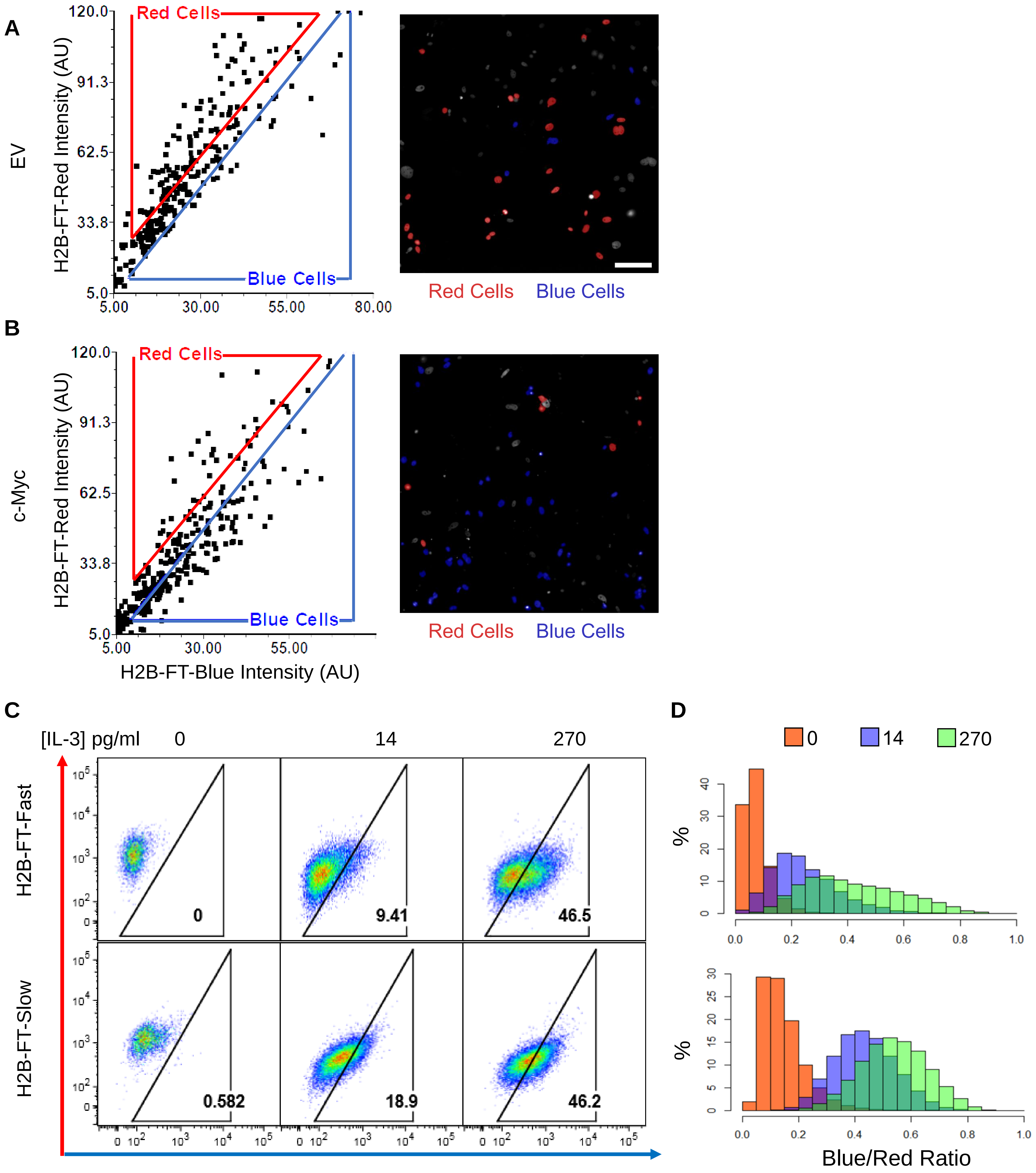


**Figure S3. H2B-FT blue/red ratio reflects proliferative changes induced by oncogenes or cytokine concentration. A-B.** Image cytometry of MEFs transduced with either an empty control (A) or c-Myc overexpression vector (B) were captured by time-lapse microscopy for several days. The raw images were subjected to a quantitative processing workflow. Nuclear measurements of integrated red vs. blue fluorescence intensity were plotted for individual cells within each specific gate. Data from a representative well, combining four fields of view, are shown on the left. Representative fields of view of cells corresponding to the blue/red gating are shown on the right. Images are shown at the same scale. Scale bar = 100µm. **C.** Representative FACS plots of BaF3 cells expressing the H2B-FT-Fast (top) and H2B-FT-Slow (bottom) mutational variants, grown under different IL-3 concentrations. **D.** Flow cytometry data from (C) plotted as histograms of blue/red ratio.

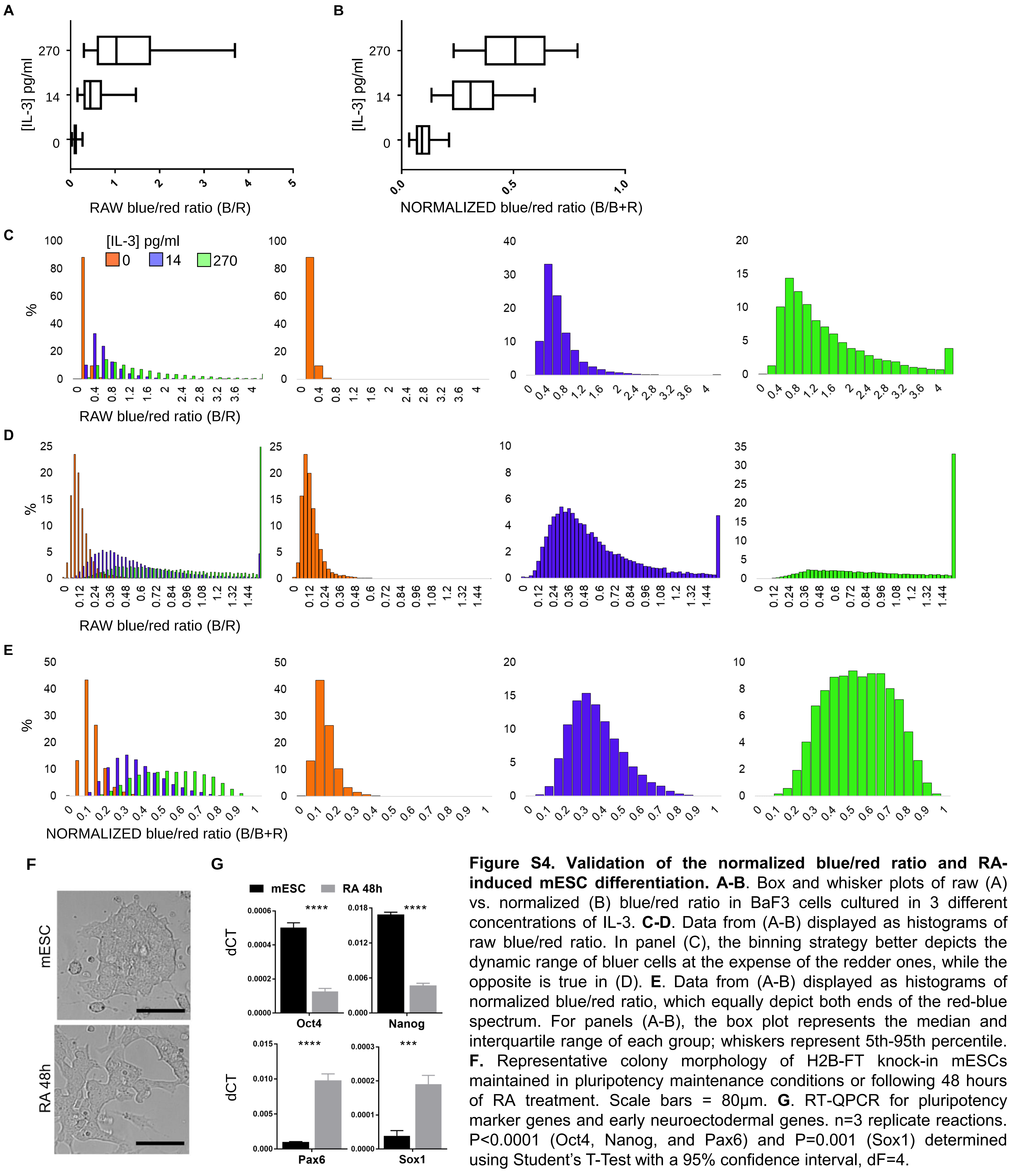


**Figure S4. Validation of the normalized blue/red ratio and RA-induced mESC differentiation. A-B**. Box and whisker plots of raw (A) vs. normalized (B) blue/red ratio in BaF3 cells cultured in 3 different concentrations of IL-3. **C-D**. Data from (A-B) displayed as histograms of raw blue/red ratio. In panel (C), the binning strategy better depicts the dynamic range of bluer cells at the expense of the redder ones, while the opposite is true in (D). **E**. Data from (A-B) displayed as histograms of normalized blue/red ratio, which equally depict both ends of the red-blue spectrum. For panels (A-B), the box plot represents the median and interquartile range of each group; whiskers represent 5th-95th percentile. **F.** Representative colony morphology of H2B-FT knock-in mESCs maintained in pluripotency maintenance conditions or following 48 hours of RA treatment. Scale bars = 80µm. **G**. RT-QPCR for pluripotency marker genes and early neuroectodermal genes. n=3 replicate reactions. P<0.0001 (Oct4, Nanog, and Pax6) and P=0.001 (Sox1) determined using Student's T-Test with a 95% confidence interval, dF=4.

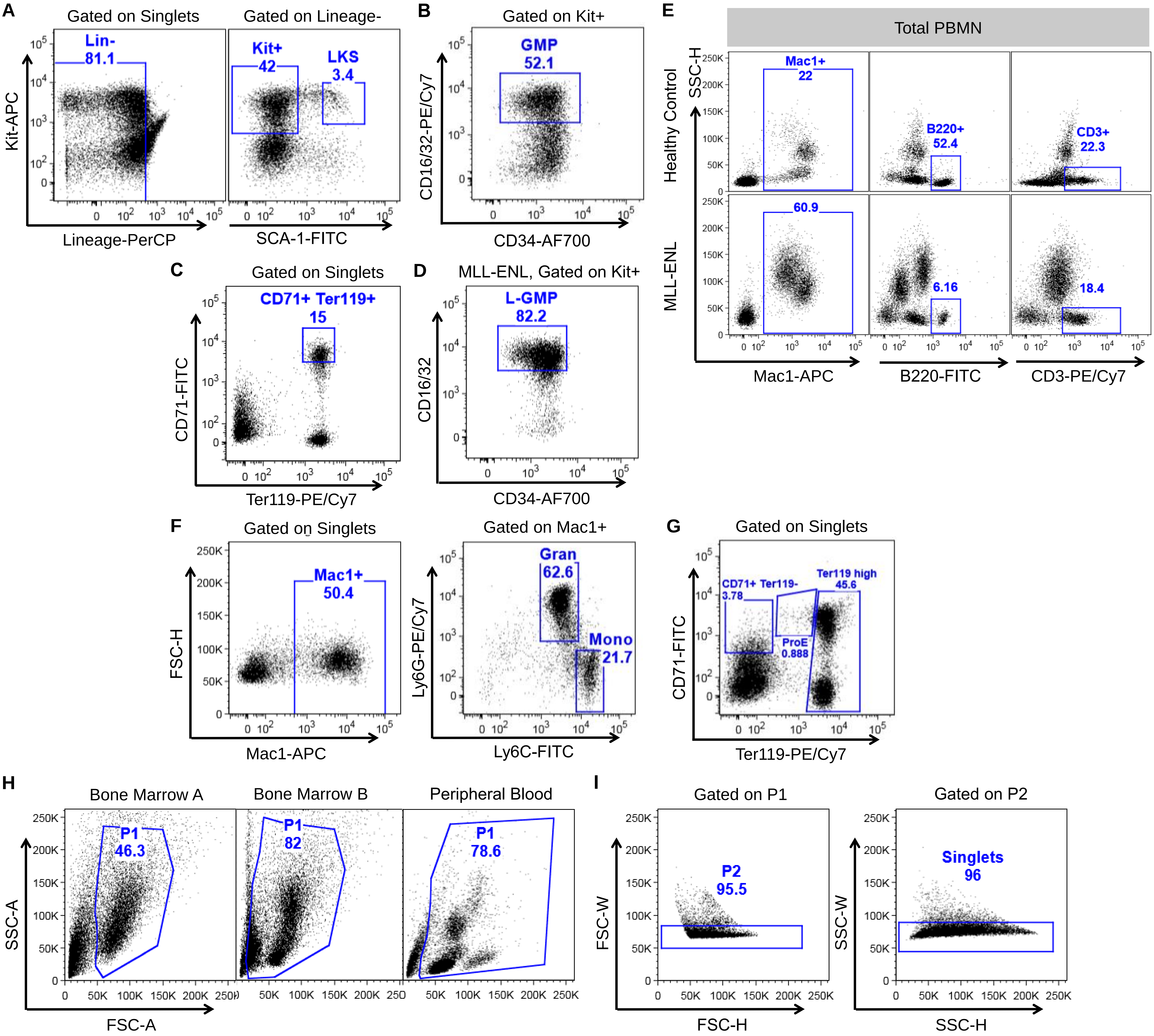


**Figure S5. Identification of specific hematopoietic populations in bone marrow and peripheral blood. A-B.** Gating for LKS and GMP. **C.** Gating for CD71+/Ter119+ erythroid cells. **D.** Gating for L-GMP compartment from H2B-FT x iMLL-ENL mice following 8 days of Dox treatment. **E.** Myeloid (Mac1+), B-cell (B220+), and T-cell (CD3+) compartments in peripheral blood mononuclear (PBMN) cells of healthy H2B-FT mice (top) and H2B-FT x iMLL-ENL mice following 8 days of Dox treatment (bottom). **F.** Gating for bone marrow monocytic and granulocytic cells. **G.** Gating strategy for distinct stages of erythroid differentiation. **H.** Initial determination of live cells by forward- and side-scatter gating. All bone marrow populations (A-D, F) were gated as shown in “Bone Marrow A”, except for erythroid cells (G) which were gated as shown in “Bone Marrow B”. PBMN (E) were gated as shown in “Peripheral Blood”. **I.** Doublet exclusion strategy used for all bone marrow and PBMN populations.

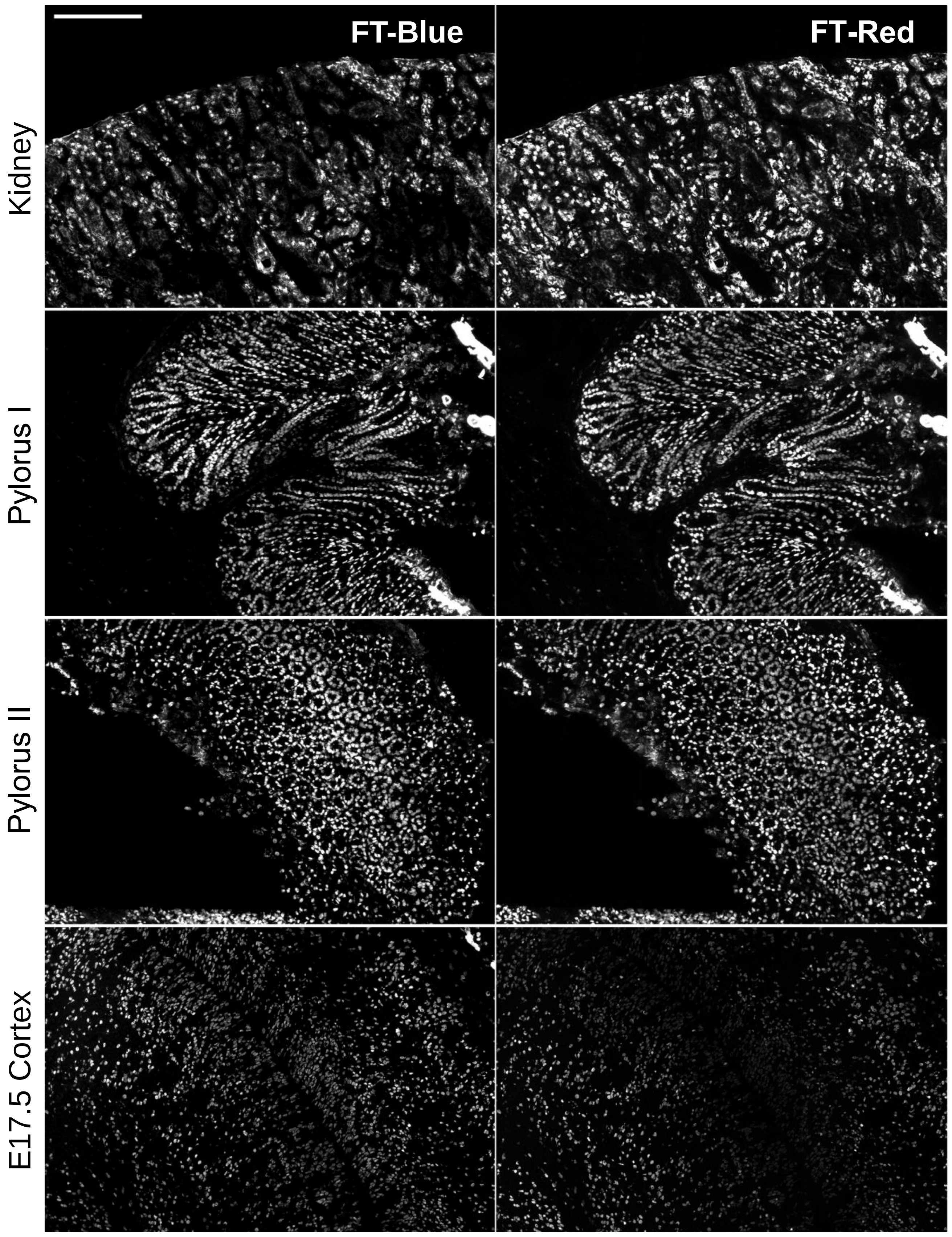


**Figure S6. H2B-FT blue/red ratio is consistent with relative turnover rates in solid tissue sections.** Single-channel fluorescence images showing H2B-FT-Blue and H2B-FT-Red in solid tissue sections. Kidney (top) and pylorus in two orientations (middle) from a representative adult iH2B-FT mouse are shown. Bottom, the neocortex of a representative E17.5 iH2B-FT mouse embryo. Scale bar = 200µm.

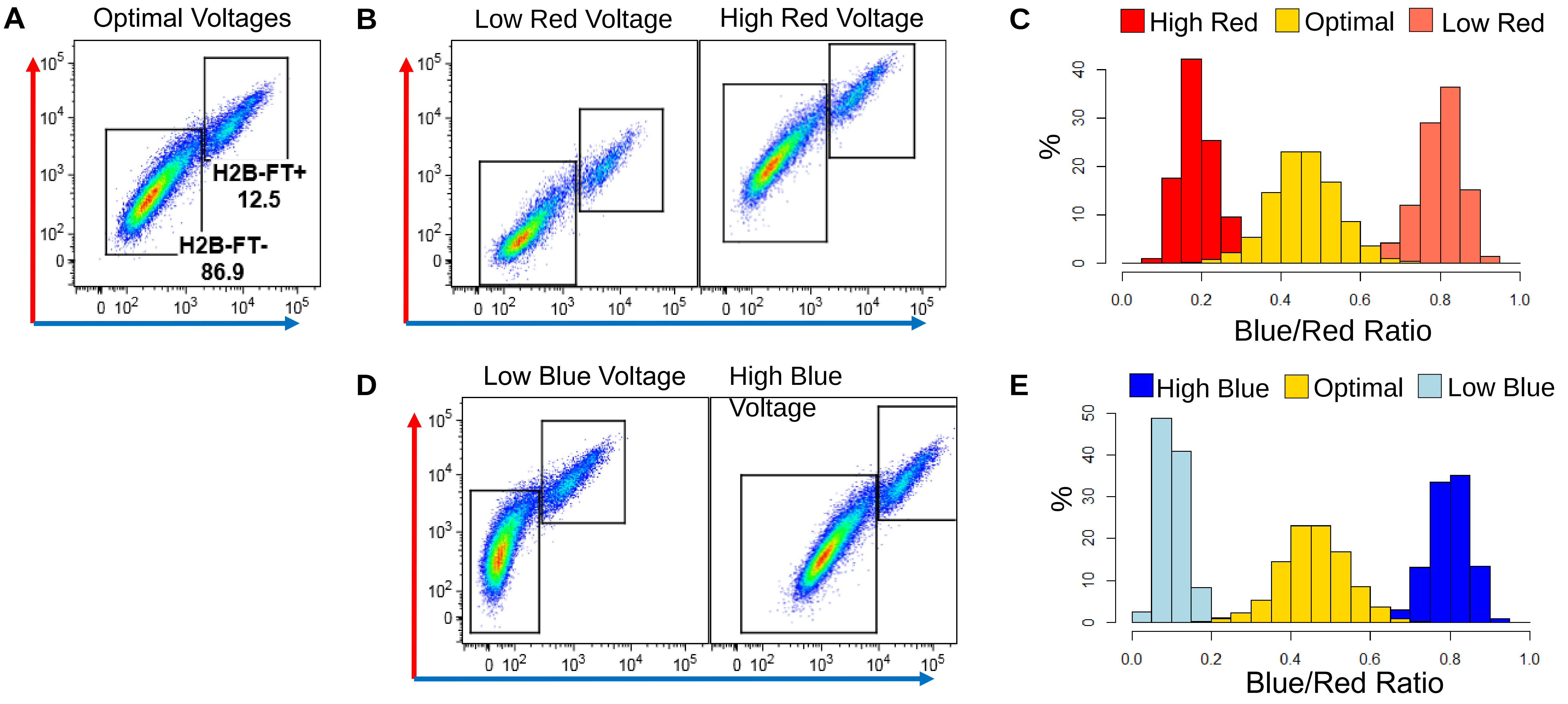


**F**

Equation S2b:

$$\frac{\bar{B}}{\bar{R}} = \frac{C \cdot \tau_C}{\tau_D}$$

Equation S2b, Normalized:

$$\frac{\bar{B}}{\bar{B} + \bar{R}} = \frac{C \cdot \tau_C}{C \cdot \tau_C + \tau_D}$$

**G**

| CC length percentile | Actual CC length (h) | Blue/red ratio percentile | Blue/red ratio value | Derived $C \cdot \tau_C$ Value | Prediction 1: CC length (h) | Prediction 2: CC length (h) | Prediction 3: CC length (h) | Prediction 4: CC length (h) |
|---|---|---|---|---|---|---|---|---|
| 10 | 9.12 | 90 | 0.711 | **22.44** | **9.12** | 8.19 | 7.95 | 7.06 |
| 25 | 10.7 | 75 | 0.653 | **20.14** | 11.92 | **10.70** | 10.40 | 9.23 |
| 50 | 14 | 50 | 0.583 | **19.57** | 16.05 | 14.41 | **14.00** | 12.42 |
| 80 | 21.4 | 20 | 0.448 | **17.37** | 27.65 | 24.82 | 24.11 | **21.40** |

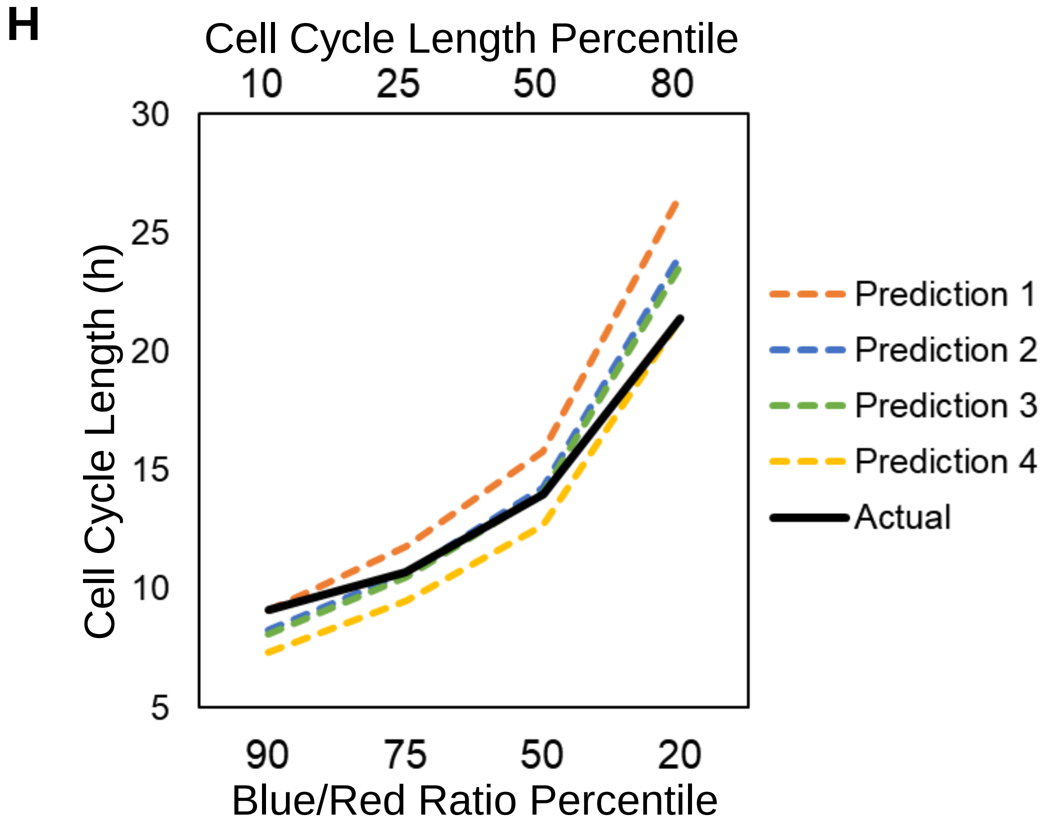


**I**

Equation S2d:

$$\frac{\bar{B}}{\bar{R}} = C \cdot \tau_C \left(\frac{1}{\tau_D} + \frac{1}{\tau_R}\right)$$

Equation S2d, Normalized:

$$\frac{\bar{B}}{\bar{B} + \bar{R}} = \frac{C \cdot \tau_C}{C \cdot \tau_C + (\frac{1}{\tau_D} + \frac{1}{\tau_R})^{-1}}$$

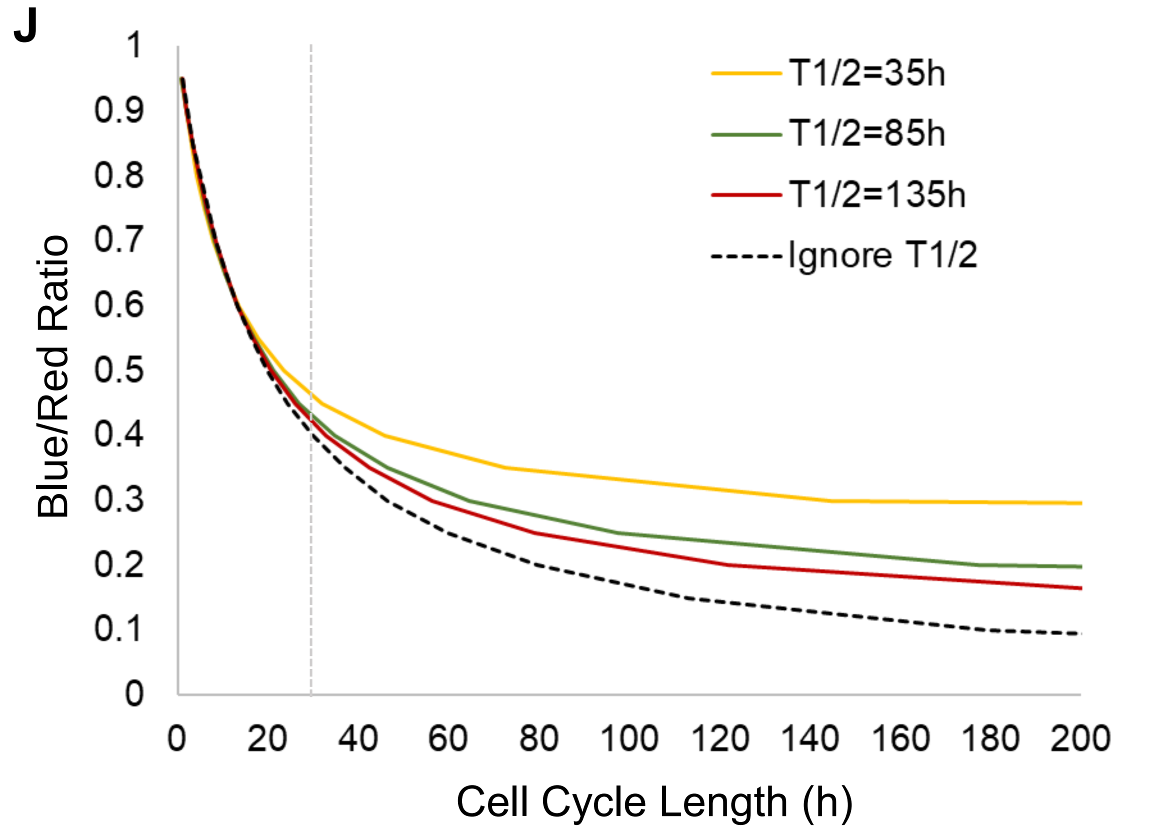


**K**

| Blue/Red Ratio | Calculated CC Length (h) Given by Red $T_{1/2}$ = 35h | Calculated CC Length (h) Given by Red $T_{1/2}$ = 85h | Calculated CC Length (h) Given by Red $T_{1/2}$ = 135h | CC Length, Mean of 3 Calculations | Error Range (h) | Error Range as % of Mean Prediction |
|---|---|---|---|---|---|---|
| 0.25 | 1013.91 | 97.61 | 79.09 | 396.87 | 934.8 | 235.5 |
| 0.3 | 144.37 | 64.51 | 56.42 | 88.43 | 88.0 | 99.5 |
| 0.35 | 72.57 | 46.37 | 42.40 | 53.78 | 30.2 | 56.1 |
| 0.4 | 45.92 | 34.91 | 32.87 | 37.90 | 13.1 | 34.4 |
| 0.45 | 32.02 | 27.02 | 25.97 | 28.34 | 6.1 | 21.4 |
| 0.5 | 23.49 | 21.26 | 20.75 | 21.83 | 2.7 | 12.6 |
| 0.55 | 17.72 | 16.86 | 16.65 | 17.08 | 1.1 | 6.3 |
| 0.6 | 13.56 | 13.40 | 13.36 | 13.44 | 0.2 | 1.5 |
| 0.65 | 10.41 | 10.60 | 10.65 | 10.55 | 0.2 | 2.2 |
| 0.7 | 7.95 | 8.29 | 8.38 | 8.21 | 0.4 | 5.2 |
| 0.75 | 5.98 | 6.35 | 6.46 | 6.26 | 0.5 | 7.7 |
| 0.8 | 4.35 | 4.70 | 4.80 | 4.62 | 0.4 | 9.7 |
| 0.85 | 3.00 | 3.28 | 3.37 | 3.22 | 0.4 | 11.5 |
| 0.9 | 1.85 | 2.05 | 2.11 | 2.00 | 0.3 | 13.0 |
| 0.95 | 0.86 | 0.96 | 0.99 | 0.94 | 0.1 | 14.3 |

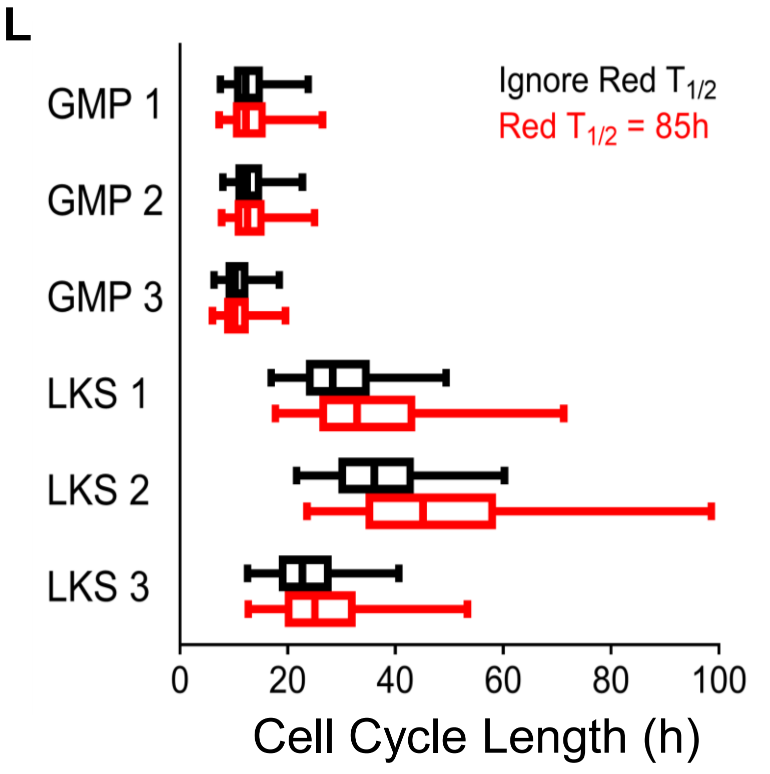


**Figure S7. Determining critical parameters for calculating cell cycle length from blue/red ratio.** (Legend on following page)

**Figure S7. Determining critical parameters for calculating cell cycle length from blue/red ratio. A.** Flow cytometry plot showing the red vs. blue color profile of primary MEFs expressing H2B-FT. “Optimal” voltage settings were carefully determined using single-color controls and all-negative controls. Altering fluorescence detection parameters impacts the dynamic range of blue/red ratio as follows: **B.** The same sample as in (A) analyzed using lower (left) or higher (right) detection voltages for the red fluorescence. **C.** Flow cytometry data from (A-B) plotted as histograms of blue/red ratio. **D.** The same cells as in (A-C) analyzed using lower (left) or higher (right) detection voltages for the blue fluorescence. **E.** Flow cytometry data from (A) and (D) plotted as histograms of blue/red ratio. **F**. The mathematical model detailed in Equation S2 predicts that cell cycle length can be determined from the steady-state blue/red ratio of cells expressing H2B-FT. Since red molecules are stable, the rate of red removal ($\delta$) can be simplified to be inversely proportional to the cell cycle time ($\tau_D$). $\tau_C$ is the blue-to-red conversion time and $C$ is a normalization constant. The equation can also be solved using the normalized blue/red ratio value, B/(B+R). **G.** Four paired data points of corresponding cell cycle (CC) length percentile and blue/red ratio percentile (as shown in Figure 7A-B) were used to derive the $C \cdot \tau_C$ value using the equation in (F). Each $C \cdot \tau_C$ value (shown in bold) was then applied onto the other three blue/red ratios to generate four separate predictions of cell cycle lengths. Bold cell cycle lengths indicate the experimentally observed value. **H.** The four predicted cell cycle length curves were compared with the actual data (solid black line). From this plot, a $C \cdot \tau_C$ value representing the average of predictions 2 and 3 was selected for estimating *in vivo* cell cycle lengths. **I**. The equation was modified to incorporate a rate constant, $\tau_R$, for active protein degradation. $\tau_R$ is related to protein half-life via the formula $T_{1/2} = \tau_R . \ln 2$. **J**. The effect of red half-life on the quantitative model relating blue/red ratio and cell cycle length. Since the value of the H2B-FT-Red half-life *in vivo* is currently unknown, we derived curves using the formula in (I) by inputting various half-life values ranging from 35-135 hours. Black lines in each plot show the curve obtained using the simplified version of Equation S2 (F), in which red decay is ignored. Vertical dotted gray line shows the distance between the predictions at cell cycle length = 30h. **K.** Table showing the expected error range in cell cycle length calculations if the H2B-FT-Red half-life varied from 35h (similar to that in Hela cells, see Figure S2D), to 85h (similar to that in MEFs, see Figure S2C) and to 135h. Yellow box indicates the approximate cell cycle length at which the H2B-FT reporter read-out is predicted to be least sensitive to variable rates of H2B-FT-Red degradation. **L.** Cell cycle length distributions of LKS and GMP populations from three mice (as in Figure 7F) calculated using different red fluorescent half-life parameters. Boxes represent the median and interquartile range of each group; whiskers represent 5th-95th percentile.

**Movie S1. Time lapse of H2B-FT in HeLa cells following Dox-induced expression.** Time stamp format is hh:mm.

**Movie S2. Time lapse of H2B-FT in HeLa cells with constitutive H2B-FT expression**. Time stamp format is hh:mm.

| **Cloning** | **Sequence (5'-3')** |
|---|---|
| Gateway *FT-Fast/Slow* Fwd | GGGGACAACTTTGTACAAAAAAGTTGCCACCATGGTGAGCAAGGGCGAG |
| Gateway *FT-Medium* Fwd | GGGGACAACTTTGTACAAAAAAGTTGCCACCATGGTAAGCAAGGGCGAG |
| Gateway *FT-Fast/Slow* Rev | GGGGACAACTTTGTACAAGAAAGTTGGCAATTACTTGTACAGCTCGTCCATG |
| Gateway *H2B* Fwd | GGGGACAACTTTGTACAAAAAAGTTGGCACCATGCCAGAGCCAGCGAAGTCT |
| *H2B-FT-Fast/Slow* Junction Fwd | AGCGCTAAGGATCCGATGGTGAGCAAGGGCGAGGAGGATAA |
| *H2B-FT-Fast/Slow* Junction Rev | ACCATCGGATCCTTAGCGCTGGTGTACTTGGTGACGGCCTTA |
| *H2B-FT-Medium* Junction Fwd | AGCGCTAAGGATCCGATGGTAAGCAAGGGCGAGGAGGATAA |
| pSCMV *FT-Fast/Slow* Fwd | TATTAAGCTTGCCACCATGGTGAGCAAGGGCGAGGA |
| pSCMV *FT-Medium* Fwd | TATTAAGCTTGCCACCATGGTAAGCAAGGGCGAGGA |
| pSCMV *FT-Fast/Medium/Slow* Rev | CCGTCTCGAGTTACTTGTACAGCTCGTCCATGCC |
| pSCMV and p2lox *H2B* Fwd | TATTAAGCTTGGCACCATGCCAGAGCCAGCGAA |
| p2lox and pT3 *FT-Fast/Medium/Slow* Rev | CCGTGCGGCCGCTTACTTGTACAGCTCGTCCAT |
| pT3 *MCS* Fwd | CGCGTGGCGCGCCTTAATTAAGTTTAAACGC |
| pT3 *MCS* Rev | GGCCGCGTTTAAACTTAATTAAGGCGCGCCA |
| pT3 *H2B* Fwd | GTATTGTTTAAACGGCACCATGCCAGAGCCAGCGAA |
| **Genotyping** | |
| *HPRT::H2B-FT* Knock-in Fwd | CTAGATCTCGAAGGATCTGGAG |
| *HPRT::H2B-FT* Knock-in Rev | ATACTTTCTCGGCAGGAGCA |
| *HPRT::H2B-FT* WT Fwd | GTCATAGGAACTGCGGTCGT |
| *HPRT::H2B-FT* WT Rev | GCTGGGATTTGAACTCAGGA |
| *Rosa26::rtTA* WT Fwd | AAAGTCGCTCTGAGTTGTTAT |
| *Rosa26::rtTA rTTA* FWD | GCGAAGAGTTTGTCCTCAACC |
| *Rosa26::rtTA Rosa26* Rev | GGAGCGGGAGAAATGGATATG |
| *Col1::MLL-ENL Col1* Fwd | TCCCTCACTTCTCATCCAGATATT |
| *Col1::MLL-ENL* Knock-in Rev | GGACAGGATAAGTATGACATCATCAA |
| *Col1::MLL-ENL* WT Rev | AGTCTTGGATACTCCGTGACCATA |
| **qPCR** | |
| *Oct4* Fwd | TCTTTCCACCAGGCCCCCGGCTC |
| *Oct4* Rev | TGCGGGCGGACATGGGGAGATCC |
| *Nanog* Fwd | AAATCCCTTCCCTCGCCATC |
| *Nanog* Rev | TTTGGGACTGGTAGAAGAATCAGG |
| *Pax6* Fwd | ACCAGTGTCTACCAGCCAATCC |
| *Pax6* Rev | GCACGAGTATGAGGAGGTCTGA |
| *Sox1* Fwd | GCCGAGTGGAAGGTCATGTC |
| *Sox1* Rev | TTGAGCAGCGTCTTGGTCTTG |
| *GAPDH* Fwd | GGTGCTGAGTATGTCGTGGAG |
| *GAPDH* Rev | GGCGGAGATGATGACCCTTT |

**Table S1. Primer sequences used for cloning, genotyping, and qPCR.**

| Antibody | Clone | Vendor |
|---|---|---|
| Ly-6G/6C - Biotin | RB6-8C5 | BD Pharmingen™ |
| CD3e - Biotin | 145-2C11 | BD Pharmingen™ |
| CD45R/B220 - Biotin | RA3-6B2 | BD Pharmingen™ |
| CD11b - Biotin | M1/70 | BD Pharmingen™ |
| Ter119 - Biotin | Cat. 553672 | BD Pharmingen™ |
| CD8a - Biotin | 53-67 | BD Pharmingen™ |
| CD4 - Biotin | GK1.5 | BD Pharmingen™ |
| CD117 - APC | 2B8 | BD Pharmingen™ |
| Ly-6A/E (Sca-1) - FITC | E13-161.7 | BD Pharmingen™ |
| Streptavidin - PerCP | Cat. 554064 | BD Pharmingen™ |
| CD16/32 - PE-Cy7 | 93 | eBioscience |
| CD34 - AlexaFluor® 700 | RAM34 | BD Pharmingen™ |
| CD11b - APC | M1/70 | eBioscience |
| CD45R/B220 - FITC | RA3-6B2 | BD Pharmingen™ |
| CD3 - PE-Cy7 | 17A2 | BioLegend® |
| CD48 - AlexaFluor® 700 | HM48-1 | BioLegend® |
| CD150 - PE-Cy7 | TC15-12F12.2 | BioLegend® |
| CD71 - FITC | C2 | BD Pharmingen™ |
| Ter119 - PE-Cy7 | Cat. 25-5921-81 | eBioscience |
| Ly6C - FITC | AL-21 | BD Pharmingen™ |
| Ly6C - PE-Cy7 | 1A8 | BD Pharmingen™ |

**Table S2. Mouse antibodies used for FACS/flow cytometry.**

| | Red | Middle | Blue | Bulk |
|---|---|---|---|---|
| Red | | 0.1813 | 0.0003 (***) | 0.0007 (***) |
| Middle | <0.0001 (****) | | 0.0005 (***) | 0.0019 (**) |
| Blue | <0.0001 (****) | 0.0026 (**) | | 0.0188 (*) |
| Bulk | <0.0001 (****) | 0.0013 (**) | <0.0001 (****) | |

| |
|---|
| mESC |
| RA 48h |

**Table S3. Exact P-values for all comparisons in Fig. 4k.**
